# SUMOylation of MFF is required for stress-induced mitochondrial fission

**DOI:** 10.1101/2023.05.05.539603

**Authors:** Richard Seager, Nitheyaa Shree Ramesh, Stephen Cross, Chun Guo, Kevin A. Wilkinson, Jeremy M. Henley

## Abstract

Mitochondrial fission regulates mitochondrial morphology, function, mitophagy and apoptosis. Fission is mediated by the GTPase dynamin related protein-1 (DRP1) and its recruitment to the outer mitochondrial membrane by DRP1 receptors. Mitochondrial fission factor (MFF) is considered the major pro-fission receptor, whereas the mitochondrial dynamics proteins (MiD49/51) sequester inactive DRP1 and facilitate the MFF-DRP1 interaction by forming a trimeric DRP1-MiD-MFF complex. Here, we identify MFF as a target of poly-SUMOylation at a single residue (Lys^151^). Following bioenergetic stress, AMPK phosphorylates MFF to promote its SUMOylation, a critical step in stress-induced fragmentation. MFF SUMOylation is not required for DRP1 recruitment from the cytosol but causes a rearrangement of the trimeric fission complex to displace MiD proteins. This alleviates MiD inhibition of DRP1 to facilitate formation of a fission-competent complex. Thus, our data demonstrate that MFF SUMOylation fine-tunes the ratio of MiD to DRP1 for the dynamic control of stress-induced mitochondrial fragmentation.

## Introduction

Mitochondria form interconnected networks that undergo continuous cycles of fusion and fission to govern mitochondrial morphology and function. Under basal conditions, the overall morphology does not change, with balanced fusion and fission events (Detmer and Chan, 2007; Twig et al., 2008). Fusion is necessary to maintain mtDNA copy number, respiration capacity and membrane potential (Chen et al., 2005), and also helps to compensate for defects in mitochondrial function by fusing healthy mitochondria with sub-optimal functioning mitochondria (Nakada et al., 2001). Mitochondrial fission, on the other hand, ensures equal distribution of mitochondria during cell division (Taguchi et al., 2007), generates sufficiently small mitochondria for transport around the cell to sites of energy demand, particularly important in polarised cells such as neurons (Cagalinec et al., 2013; Lewis et al., 2018), and is also crucial for quality control, to isolate damaged mitochondria for removal by mitophagy (Twig et al., 2008; Kageyama et al., 2012, 2014).

The architecture of the mitochondrial network responds dynamically to fluctuating bioenergetic demands and cellular stress (Wai and Langer, 2016; Tilokani et al., 2018; Pernas and Scorrano, 2016). Moderate cell stress, such as nutrient deprivation, UV-C irradiation or cycloheximide exposure increase fusion (termed stress-induced mitochondrial hyperfusion; SIMH), which elongates mitochondria, promotes mitochondrial function and is cytoprotective (Gomes et al., 2011; Rambold et al., 2011; Tondera et al., 2009). Conversely, during mitochondrial dysfunction or severe cellular stress, such as oxidative stress or treatment with mitochondrial inhibitors, the mitochondrial network undergoes fragmentation (Losón et al., 2013; Qi et al., 2011; Toyama et al., 2016), which results in reduced mitochondrial function, increased mitophagy, and is associated with apoptosis (Frank et al., 2001; Kageyama et al., 2014; Leboucher et al., 2012; Losón et al., 2013). Thus, the balance of mitochondrial fusion/fission is fundamental for mitochondrial function and cellular homeostasis, and dysregulation of these systems is a prominent feature in multiple diseases (Chan, 2020).

Fission is mediated by recruitment of DRP1 from the cytosol to the mitochondrial outer membrane, where it oligomerises and powers membrane scission via GTP hydrolysis (Mears et al., 2011; Smirnova et al., 2001). There are four known DRP1 receptors: Fis1, MFF, MiD49 and MiD51, each of which can independently recruit DRP1 to the mitochondrial surface (Losón et al., 2013; Osellame et al., 2016; Palmer et al., 2013). MFF is the main pro-fission receptor and its overexpression fragments mitochondria, whereas knockdown results in severe mitochondrial elongation (Losón et al., 2013; Otera et al., 2010). Although MiD49/51 can recruit DRP1 (Otera et al., 2016; Palmer et al., 2011, 2013), their effect on fission is less clear, since their overexpression increases DRP1 recruitment, but also causes mitochondrial elongation (Palmer et al., 2011, 2013; Zhao et al., 2011), likely due to sequestering inactive DRP1 (Losón et al., 2013; Palmer et al., 2011). MiD receptors bind to a broad range of DRP1 oligomeric states, including DRP1 mutants with impaired GTPase activity or lacking the ability to form higher oligomeric states (Yu et al., 2021). In contrast, it has been reported that MFF favours binding to higher order DRP1 oligomers and has significantly impaired ability to interact with mutants lacking GTPase activity or the capacity to oligomerise (Yu et al., 2021; Liu and Chan, 2015). Although Fis1 has important roles in mitophagy and asymmetric mitochondrial division (Kleele et al., 2021; Shen et al., 2014; Waters et al., 2022), it plays a relatively minor role in fission (Otera et al., 2010; Osellame et al., 2016).

DRP1, MiD and MFF form a trimeric complex in which MiD proteins facilitate the MFF-DRP1 interaction (Yu et al., 2017). Thus, the MiD proteins serve as a platform for DRP1 recruitment and assembly, and the differential association of mitochondrially bound DRP1 with the receptors determines fission (Yu et al., 2021). However, how this trimeric complex adapts rates of mitochondrial fission to meet fluctuating bioenergetic demands and mitochondrial stress remains poorly understood.

AMP-activated protein kinase (AMPK) is a stress response kinase that maintains energy homeostasis. During times of enhanced energy expenditure (signalled by an increase in the AMP/ATP ratio), AMPK is activated to promote energy-producing processes while minimising energy-demanding processes (Herzig and Shaw, 2018). AMPK phosphorylates MFF at Ser^155^ and Ser^172^ in response to mitochondrial stress (Ducommun et al., 2015; Toyama et al., 2016), a modification that is necessary and sufficient to promote mitochondrial fission (Toyama et al., 2016) and has been shown to occur during mitophagy (Seabright et al., 2020). However, how this phosphorylation event regulates the fission machinery at a molecular level is an important unanswered question.

The post-translational modifier protein small ubiquitin-like modifier (SUMO) is reversibly conjugated to target proteins to regulate their functions, interactions and activity. Sentrin-specific proteases (SENP1-3, 5-7) regulate the deconjugation of SUMO from targets. SUMOylation generally occurs within the consensus sequence ψKxD/E, where ψ is a large hydrophobic residue, and x is any amino acid (Wilkinson and Henley, 2010; Flotho and Melchior, 2013). The major SUMO isoforms are SUMO1-3, with SUMO2 and SUMO3 forming poly-SUMO chains, often terminated by SUMO1 (Tatham et al., 2001; Matic et al., 2008). Mitochondrial protein SUMOylation regulates mitochondrial morphology and function (Zunino et al., 2007; Prudent et al., 2015). For example, SUMO1-ylation stabilises DRP1, promotes fission (Harder et al., 2004; Zunino et al., 2007) and has roles in apoptosis (Prudent et al., 2015), whereas DRP1 SUMO2/3-ylation reduces binding to MFF and inhibits cytochrome c release and cell death (Guo et al., 2017, 2013).

Here, we identify MFF as an important SUMO target, and demonstrate that AMPK-mediated phosphorylation of MFF enhances SUMOylation during times of mitochondrial stress. MFF phosphorylation/SUMOylation does not increase DRP1 binding *per se* but remodels the DRP1-MiD-MFF fission complex by displacing the inhibitory MiD receptors, thus facilitating stress-induced mitochondrial fission. These findings establish a link between MFF phosphorylation and the molecular events that govern fission, and identify MFF SUMOylation as a crucial step in coupling bioenergetic stress to dynamic regulation of mitochondrial morphology.

## Results

### MFF is SUMOylated at Lys^151^ by SUMO1 and SUMO2/3

DRP1 SUMOylation plays key roles in its recruitment to mitochondria and regulation of morphology (Guo et al., 2017, 2013; Harder et al., 2004; Prudent et al., 2015; Zunino et al., 2007). We therefore wondered whether these processes were also regulated by SUMOylation of the DRP1 receptors. To test this, we transfected HEK293T cells with FLAG-SUMO and GST-DRP1 receptors. Following pulldowns and blotting for FLAG, we observed a robust ladder of SUMO1 and SUMO2 conjugation to MFF (Fig 1A, S1A), indicative of poly-SUMO chains. No FLAG signal was detected for the other receptors, indicating MFF is likely the only DRP1 receptor SUMOylated under these conditions.

**Figure 1.**
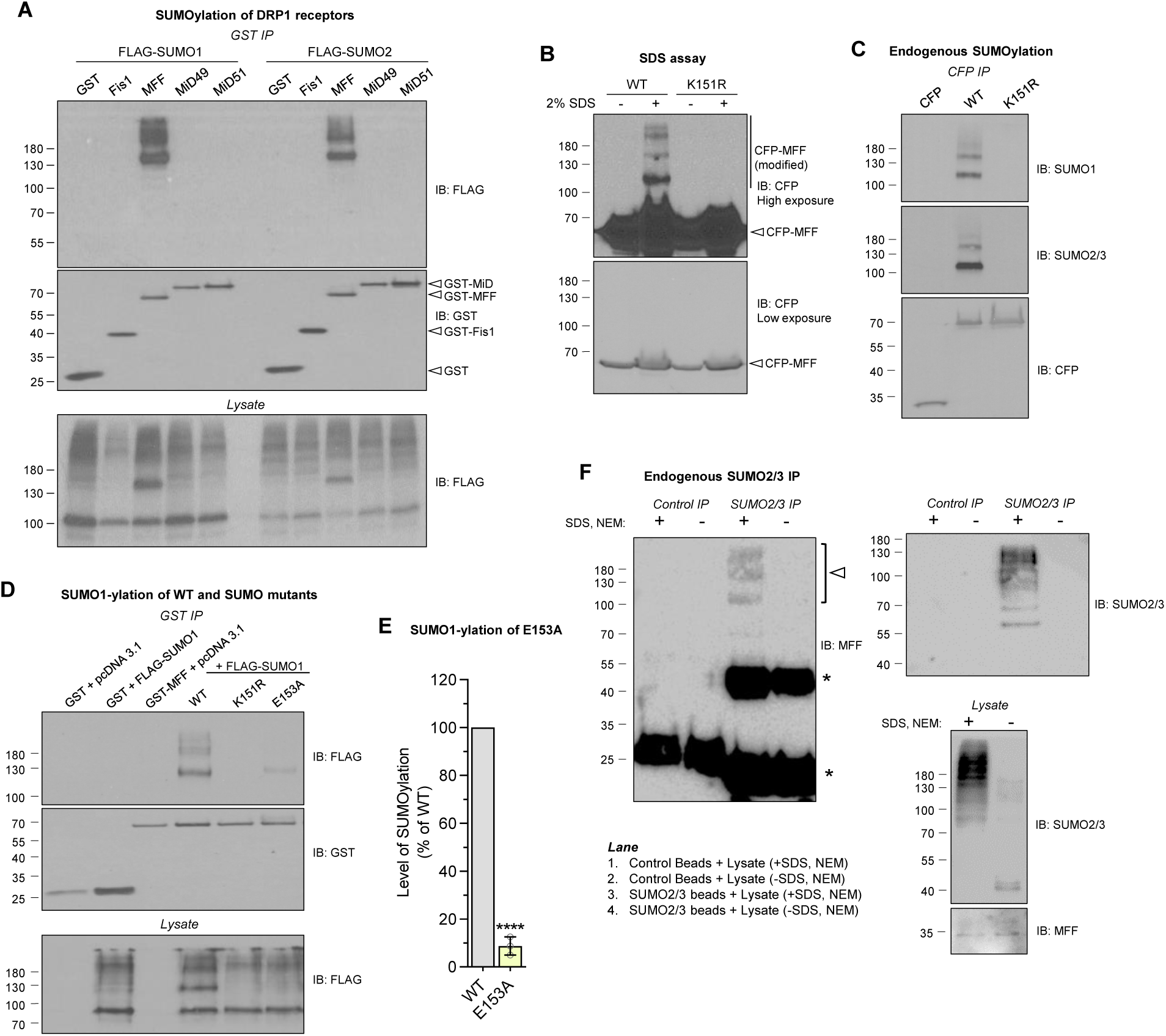
MFF is poly-SUMOylated at Lys^151^. (**A**) HEK293T cells were co-transfected with GST-tagged DRP1 receptors (Fis1 (mouse), MFF (human, isoform 1), MiD49 or MiD51 (mouse)) and either FLAG-SUMO1 or FLAG-SUMO2. GST immunoprecipitates and lysate was immunoblotted for FLAG and GST. (**B**) WT or K151R CFP-MFF transfected cells were lysed in buffer ±2% SDS and probed for CFP. (**C**) Blot of endogenously SUMOylated MFF. CFP-MFF (WT or K151R) immunoprecipitates from HEK293T cells were probed for endogenous SUMO1 or SUMO2/3. (**D**) Analysis of MFF SUMOylation-deficient mutants. GST pulldowns of the indicated mutants were blotted for FLAG and GST. (**E**) Quantification of SUMO-deficient MFF mutants, showing normalised FLAG signal to GST, expressed as percentage of WT ±S.D, n=3, p***<0.001, one-sample t-test. (**F**) SUMO2/3 IP from HEK293T cell lysate, probed for MFF. Lanes 1-2 are control lanes using protein G beads. Lanes 3-4 are SUMO2/3 enriched samples using anti-SUMO2/3 conjugated beads. HEK293T cells were lysed in buffer ±4% SDS and 20mM NEM, to preserve or inhibit SENP activity in the lysate. 4% SDS was then diluted to 0.1% in lysis buffer before incubation with beads. Enrichment of SUMO2/3 conjugated proteins in lane 3 was confirmed by SUMO2/3 reprobe (top right blot), and deconjugation of SUMO2/3 confirmed in the lysate blot. Arrow indicates endogenous SUMOylated MFF, asterisk indicates non-specific antibody bands.

The highly conserved sequence ^150^LKRE^153^ (corresponding to Lys^151^ in human MFF isoform 1 and Lys^125^ in isoforms 2-5) conforms to the SUMO consensus motif and was identified as a candidate SUMOylation site in MFF (Fig S1B-C). Lys^151^ was mutated to a non-SUMOylatable arginine (K151R) and MFF SUMOylation investigated by expressing WT or K151R CFP-MFF in HEK293T cells followed by blotting for CFP after lysis under strong denaturing conditions (±2% SDS, to retain SUMO conjugation by impairing the activity of deSUMOylating enzymes). The band-shifted modified forms of CFP-MFF disappear for MFF WT when SDS is absent and are not present in either condition with the MFF K151R mutant (Fig 1B). We then probed CFP-MFF WT and K151R immunoprecipitates for endogenous SUMO1 or SUMO2/3 and observed a complete absence of SUMOylation in the K151R mutant (Fig 1C). Finally, mutation of the adjacent E153 residue in CFP-MFF, to disrupt the SUMO consensus sequence while retaining the modifiable lysine, also results in a severe decrease in SUMOylation (Fig 1D, E). Together, these experiments identify Lys^151^ as the sole SUMO site in MFF.

We next validated SUMOylation of endogenous MFF. To do this, we used SUMO2/3 antibody-conjugated beads to immunoprecipitate endogenous SUMO2/3 conjugates from HEK293 cells, followed by immunoblotting for MFF. As a negative control we lysed cells in the absence of SDS and NEM, to retain endogenous SENP activity in the lysate to facilitate deSUMOylation. As shown in figure 1F, SUMOylated proteins were enriched in the SUMO2/3 immunoprecipitates, and a MFF immunoreactive ladder of high molecular weight species, similar in pattern to those observed in figure 1A-D, was detected only in the SUMO2/3 IP samples lysed in the presence of SDS/NEM. This MFF immunoreactive band was absent when cells were lysed in the absence of SDS and NEM, confirming MFF is endogenously SUMOylated (Fig 1F).

We have previously reported that MFF is polyubiquitinated (Lee et al., 2019). To dissect the nature of poly-SUMOylation on MFF, and to determine if mixed SUMO and ubiquitin chains are present, we performed an *in vitro* deSUMOylation and deubiquitination assay of WT CFP-MFF immunoprecipitated from HEK293T cells, using recombinant SENP1 or USP2, respectively. SENP treatment removed SUMO, but had no effect on ubiquitin, and USP2 treatment removed ubiquitin from MFF without removing SUMO. These data indicate that these two modifications are independent and that there are no mixed SUMO-ubiquitin chains on MFF (Fig S1D).

### MFF SUMOylation is not required for basal DRP1 recruitment or morphology

As reported previously (Losón et al., 2013), MFF-KO MEF cells (lacking all isoforms of MFF) exhibit an elongated and fused mitochondrial network (Fig 2A, B). We performed colocalisation analysis of DRP1 with mitochondria and observed a decrease in mitochondrially-associated DRP1 in MFF-KO cells (Fig 2C). Moreover, quantification of the mitochondrial network showed an increase in the number of branches and average mitochondrial length in the MFF-KO cells (Fig 2D, E) and a reduction in the free-end index (a parameter to determine the extent of fragmentation; Fig 2F).

**Figure 2.**
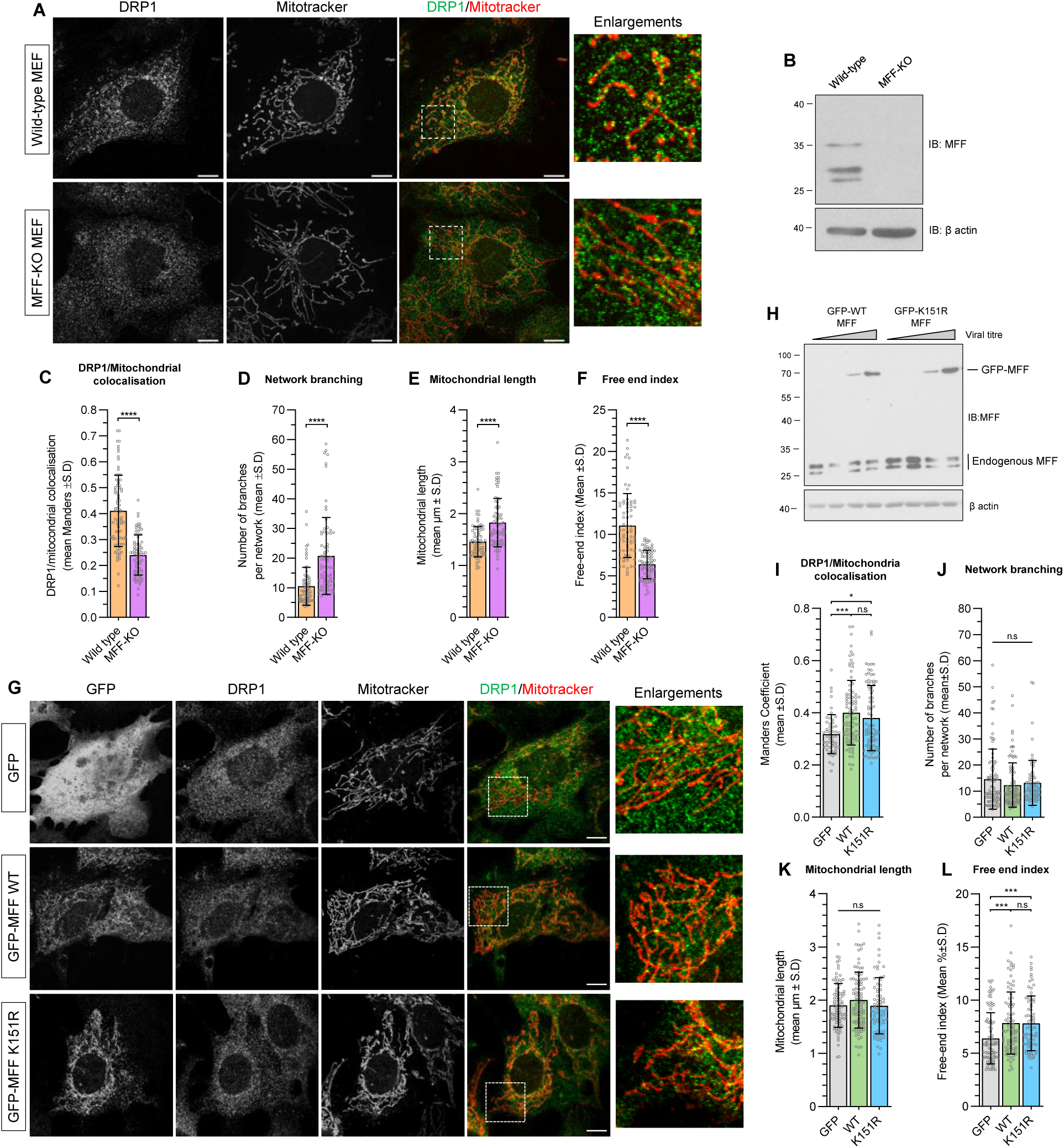
Mitochondrial morphology and DRP1 recruitment in MFF-KO MEF cells expressing MFF-WT or MFF-K151R. (**A**) Confocal images of wild-type and MFF-KO MEF cells stained for DRP1 and mitochondria using mitotracker. Enlargements show zoomed section of the highlighted area. Scale bar 10µm. (**B**) MEF cell lysate probed for MFF. MFF-KO cells lack all detectable isoforms of MFF. (**C**) Manders’ colocalisation quantification of DRP1 and mitotracker, n=3, 84-98 cells imaged. (**D-F**) Mitochondrial morphology analysis of MEF wild-type and MFF-KO cells. (**D**) Mean number of branches per network, (**E**) mean mitochondrial length and (**F**) mean free-end index. Mann-Witney test used to determine significance, 70-74 cells imaged from three independent experiments, p****<0.0001. (**G**) Confocal images of MFF-KO MEF cells expressing either GFP, GFP-MFF-WT or MFF-K151R. Enlargements show zoomed section of the highlighted area. Scale bar 10µm. (**H**) Viral titres of GFP, GFP-MFF WT and K151R infection of wild-type MEF cells were used to determine appropriate viral amount to infect cells with. The volume in lanes 4 and 8 were used for subsequent experiments. (**I**) Manders’ colocalisation of DRP1 and mitotracker (GFP, n=2, 52 cells; GFP-MFF (WT and K151R), n=3, 85-91 cells) (**J-L**) Mitochondrial morphology analysis of MFF-KO cells expressing GFP, WT-MFF or K151R-MFF. (**J**) Network branching, (**K**) mitochondrial length and (**L**) free-end index. n=3, 73-95 cells, p*<0.05, p***<0.0005, Kruskal-Wallis test followed by Dunn’s multiple comparison test.

To define the role of MFF SUMOylation in basal mitochondrial morphology, we used lentivirus to express GFP-tagged MFF-WT or K151R in MFF-KO cells, labelled mitochondria with mitotracker and stained for DRP1 (Fig 2G). Viral titres were adjusted to ensure expression of MFF constructs at approximately endogenous levels (Fig 2H). Note lanes 4 and 8 express at similar levels to each other and to endogenous MFF, so these titres were used in all subsequent experiments. Both MFF-WT and K151R localised to mitochondria and rescued DRP1 recruitment to a similar extent (Manders’ values = 0.40 and 0.38, respectively, Fig 2I), to levels similar to the wild-type MEF cells (Manders’ value = 0.41, Fig 2C). Quantitative analysis of mitochondrial morphology showed no differences in network branching (Fig 2J) or mitochondrial length (Fig 2K) between GFP, GFP-MFF WT or K151R.

We did, however, detect an increase in the free-end index for both WT and K151R-MFF above the GFP control, but there was no difference between WT and non-SUMOylatable MFF (6.4% (GFP) vs 7.8% (MFF-WT and K151R), Fig 2L). These data indicate that viral expression of WT or K151R-MFF in MFF-KO cells can at least partially rescue the fission defects, and indicate that MFF SUMOylation is not required for basal DRP1 recruitment. Furthermore, preventing MFF SUMOylation had no discernible effects on mitochondrial architecture under basal conditions.

### Phosphorylation at Ser^155^ and Ser^172^ promotes MFF SUMOylation

In response to bioenergetic stress AMPK phosphorylates MFF at Ser^155^ and Ser^172^, leading to mitochondrial fragmentation (Toyama et al., 2016; Seabright et al., 2020; Ducommun et al., 2015). Ser^155^ lies within a phosphorylation-dependent SUMO consensus motif (PDSM: Ψ-K-x-E-x-(x)-S, Fig S1C), a strong predictor of positive regulation of SUMOylation by phosphorylation (Hietakangas et al., 2006). We therefore co-expressed either double phospho-null (S155A/S172A) MFF-2SA or phospho-mimetic (S155D/S172D) MFF-2SD mutants with FLAG-SUMO1 or FLAG-SUMO2 in HEK293T cells and assessed their SUMOylation. MFF-2SD was significantly more SUMOylated than MFF-2SA (Fig 3A-D), with the 2SA mutant exhibiting significantly less SUMOylation than WT MFF. Similar results were observed for individual analysis of both the mono- and poly-SUMOylated forms of MFF (Fig S2A-D).

**Figure 3.**
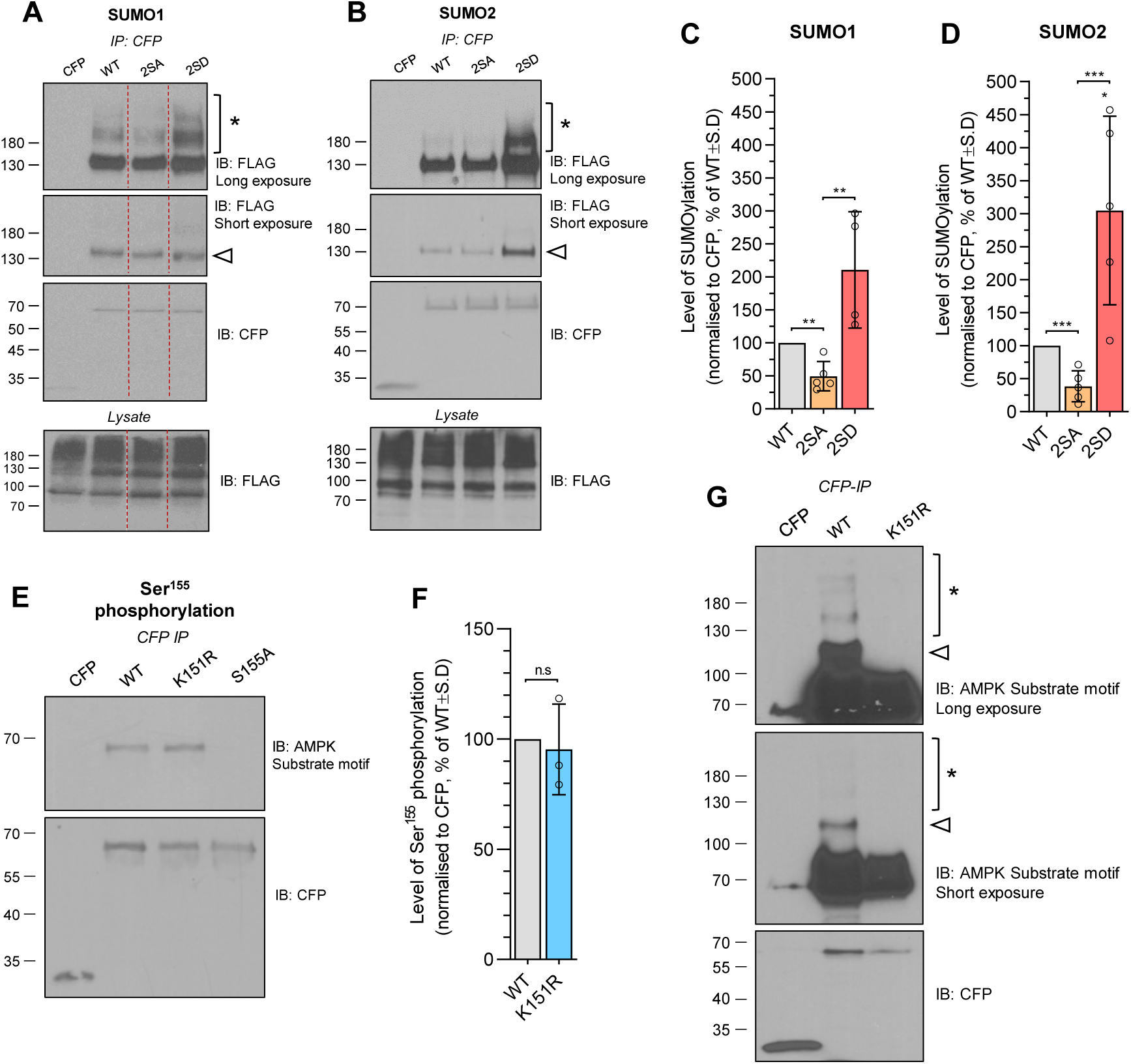
MFF SUMOylation is promoted by AMPK-mediated phosphorylation. **(A-B)** SUMOylation of MFF phosphorylation mutants. HEK293T cells were co-transfected with CFP-tagged 2SA/D MFF mutants and either **(A)** FLAG-SUMO1 or **(B)** FLAG-SUMO2. Immunoprecipitates and lysates were blotted for FLAG and CFP. Blot cropped from larger blot (Fig S2E). Arrow indicates mono-SUMOylated MFF band, asterisk shows higher molecular weight bands. **(C-D)** Quantification of SUMOylation of MFF phosphorylation mutants, expressed as mean percentage of WT ±S.D. One sample t-test performed to determine significance between mutants and WT, unpaired t-test performed to determine significance between mutants, n=4/5, p*<0.05, p**<0.01, p***<0.005. **(E)** Ser^155^ phosphorylation of MFF-K151R. HEK293T cells were transfected with the indicated CFP-MFF mutants and immunoprecipitates blotted for Ser^155^ phosphorylation using an AMPK substrate motif antibody. S155A mutant used to confirm specificity of the antibody. **(F)** Quantification of phosphorylation state of MFF-K151R, showing mean percentage of WT ±S.D. n=3, one sample t-test. **(G)** Ser^155^ phosphorylation of MFF-WT and K151R. HEK293T cells were transfected with CFP-MFF (WT or K151R) and immunoprecipitates blotted for Ser^155^ phosphorylation. Arrow corresponds to the band similar to the size of the mono-SUMOylated MFF species, asterisk represents higher molecular weight species.

To further examine the order of these two modifications, we measured Ser^155^ phosphorylation of WT and K151R-MFF using an AMPK substrate motif antibody. No signal was detected in the S155A mutant, confirming the antibody is specific for this phosphorylation site of MFF (Fig 3E). Both MFF-WT and K151R had comparable levels of Ser^155^ phosphorylation (Fig 3F), indicating that SUMOylation does not affect MFF phosphorylation. Longer exposure of phospho-Ser^155^ blots revealed a ladder of bands for MFF-WT but not for K151R (Fig 3G), similar to the SUMOylation bands observed in figure 1, suggesting these two modifications occur concurrently. Together, these data indicate that AMPK-mediated phosphorylation promotes MFF SUMOylation.

It has been reported that the mitochondrial SUMO E3 ligase MAPL, and the deSUMOylating enzymes SENP3 and SENP5, regulate DRP1 SUMOylation and mitochondrial morphology (Braschi et al., 2009; Guo et al., 2017, 2013; Prudent et al., 2015; Zunino et al., 2007). Consistent with these reports, we observed that siRNA-mediated knockdown of MAPL reduced, whereas SENP3 and SENP5 knockdown dramatically increased, MFF SUMOylation (Fig S2F-G). These findings indicate that a complex array of proteins regulate the SUMOylation status of MFF, with MAPL promoting, and SENP3/5 antagonising, MFF SUMOylation.

### DRP1 binding to MFF is independent of MFF phosphorylation and SUMOylation

AMPK-mediated phosphorylation of MFF during bioenergetic stress drives mitochondrial fission (Toyama et al., 2016). Given our data indicating that phosphorylation promotes MFF SUMOylation, we hypothesised that the high SUMOylation state of MFF-2SD would enhance DRP1 binding whereas the low SUMOylation state of MFF-2SA, or the complete lack of SUMOylation of K151R, would reduce DRP1 binding. To test this, we performed co-IP experiments of CFP-MFF mutants from transfected HEK293T cells and investigated the binding to endogenous DRP1 (Fig 4A, B, S3A). Surprisingly, there was no difference in DRP1 binding between the MFF-2SD or MFF-2SA mutants, and DRP1 binding for each of the MFF mutants was significantly reduced compared to MFF-WT (Fig 4A, B). The data for MFF-K151R and MFF-2SA are consistent with a model of MFF SUMOylation promoting fission. However, since MFF-2SD and MFF-2SA mutants have opposing effects on fission (Toyama et al., 2016), their similar binding to DRP1 was unexpected. Thus, these results indicate that there is not a simple linear sequence of MFF phosphorylation, leading to MFF SUMOylation, to promote MFF binding to DRP1.

**Figure 4.**
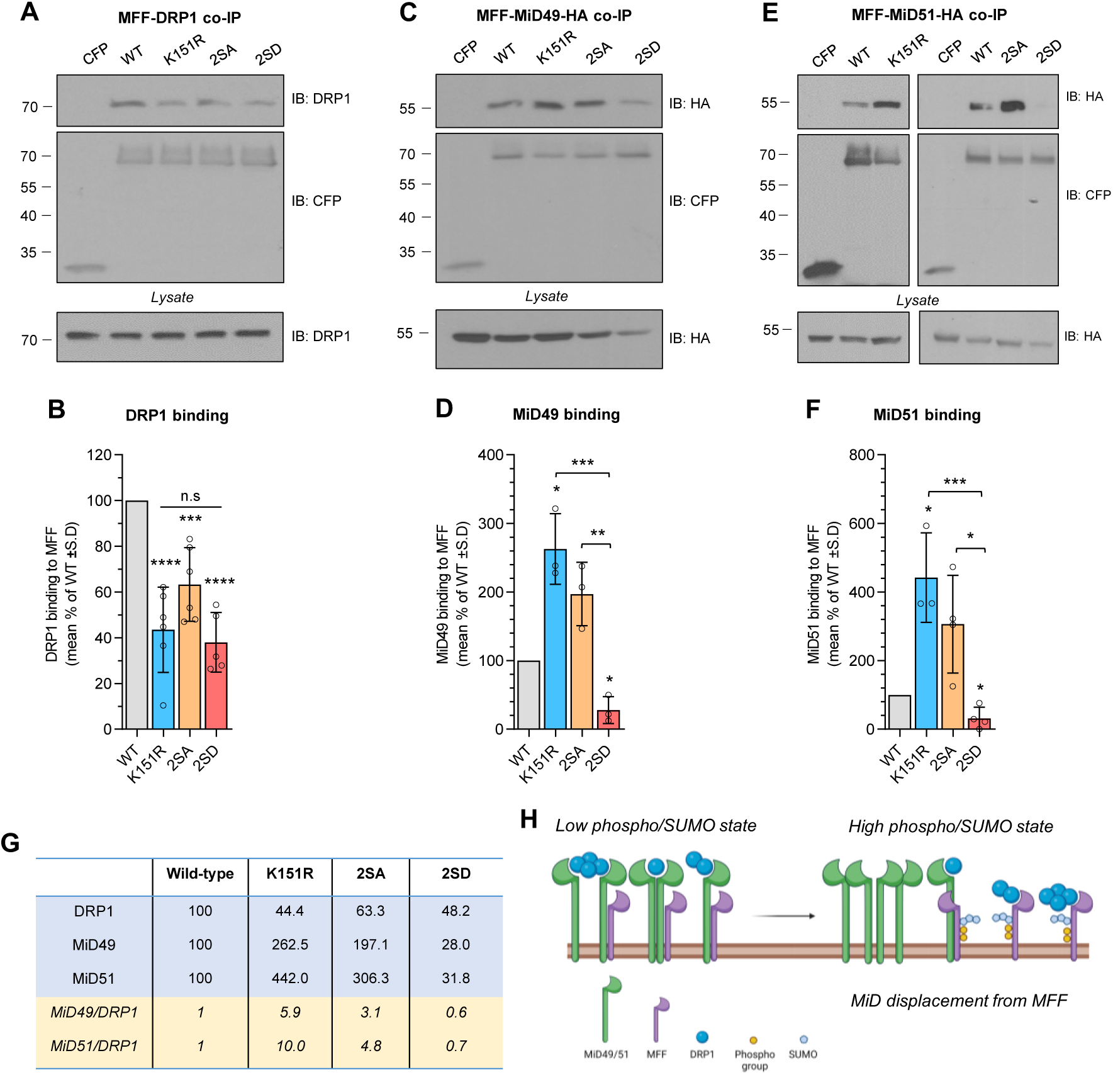
MFF post-translational modifications regulate the MiD/DRP1 ratio in the fission complex. (**A**) Representative blot of endogenous DRP1 binding to CFP-MFF mutants. HEK293T cells were transfected with the indicated CFP-MFF mutants and immunoprecipitates were probed for DRP1. (**B**) Quantification of DRP1 binding, expressed as mean percentage of WT ±S.D, n=5/6. Representative blot of (**C**) MiD49-HA and (**E**) MiD51-HA binding to MFF mutants. Immunoprecipitates from HEK293T co-transfected with MiD-HA and the indicated CFP-MFF mutants were probed for HA. (**D, F**) Quantification of (**D**) MiD49-HA and (**F**) MiD51-HA binding to MFF mutants, expressed as mean percentage of WT ±S.D, n=3/4, p*<0.05, p**<0.01, p***<0.005, p****<0.001, one-sample t-test used to determine significance from WT, one-way ANOVA used to determine significance between groups. (**G**) Table shows the relative amounts of DRP1 and MiD within the different MFF complexes, compared to WT, obtained from the quantifications (**B, D, F**). The MiD to DRP1 ratio is in italics and yellow, calculated from the values in blue. (**H**) Schematic of DRP1-MiD-MFF rearrangement in response to MFF phosphorylation and SUMOylation. Created with BioRender.com.

### SUMOylated MFF displaces MiD from the fission complex

We wondered if the similar binding of DRP1 to MFF-2SD and MFF-2SA could be explained by a mechanism in which MFF phosphorylation does not enhance DRP1 binding *per se*. Rather, we hypothesised that the stoichiometry of the DRP1-MiD-MFF trimeric fission complex (Yu et al., 2017) might be altered by MFF phosphorylation. We therefore measured MiD49-HA and MiD51-HA binding to MFF-WT, MFF-K151R, MFF-2SA and MFF-2SD under basal conditions (Fig 4C-F). Compared to MFF-WT, both MiD49 and MiD51 bound significantly more to MFF-K151R, and significantly less to MFF-2SD. Furthermore, MiD49/51-HA bound significantly more to MFF-2SA than MFF-2SD. A similar trend was also detected for endogenous MiD49 binding (Fig S3B, C).

Quantification of the relative ratios of MiD49/51 to DRP1 (Fig 4G) show that in DRP1-MiD-MFF complexes containing non-SUMOylatable MFF-K151R the ratio of MiD49 to DRP1 is ∼6-fold more than in trimeric complexes containing MFF-WT. In MFF-2SA containing complexes, the MiD49/DRP1 ratio is ∼3-fold more compared to MFF-WT, whereas the ratio for the MFF-2SD complex is ∼40% less than MFF-WT, and 5-fold less compared to MFF-2SA (Fig 4G). Likewise, the relative ratio of MiD51/DRP1 in the MFF-K151R, MFF-2SA and MFF-2SD containing complexes was 10, 4.8 and 0.7, respectively. These results demonstrate that the interplay between phosphorylation and SUMOylation of MFF modulates the stoichiometry of DRP1-MiD-MFF fission complexes by reducing levels of MiD within the assembly, rather than directly controlling MFF-DRP1 binding (Fig 4H).

DRP1 exists in multiple oligomeric states that can bind MiD49/51 (Yu et al., 2021). Since our co-IP data (Fig. 4A) do not distinguish between potentially different oligomeric states, we carried out crosslinking experiments using the non-cleavable crosslinker DSS prior to co-IP of DRP1 with MFF mutants (Fig S3D). Fission complexes containing monomeric and higher order states of DRP1 were detected, indicating that MFF-WT can bind to multiple oligomeric states of DRP1. Moreover, the MFF mutants exhibited no striking difference in binding to monomeric versus oligomeric DRP1. These results demonstrate that post-translational modifications of MFF have no discernible effect on the oligomeric states of DRP1 that MFF is capable of binding. Figure 4H is a schematic illustrating the relative differences in MiD association within the DRP1-MiD-MFF complex in response to phosphorylation and SUMOylation. This model provides an explanation of how the low phospho/SUMO state of MFF can co-IP similar levels of DRP1 to the high phospho/SUMO state, but there is less MiD in the trimeric complex containing the high phospho/SUMO state of MFF.

### CCCP-induced mitochondrial stress enhances MFF SUMOylation and displaces MiD51

We next investigated the effects of stress on MFF phosphorylation at Ser^155^ and SUMO2/3-conjugation to MFF. We first used the mitochondrial ionophore CCCP which has been used extensively to investigate mitochondrial fission (Gandre-Babbe and van der Bliek, 2008; Losón et al., 2013; Otera et al., 2010; Palmer et al., 2011) to induce fragmentation (Fig 5A). CCCP treatment caused a ∼4-fold increase in Ser^155^ phosphorylation and a concomitant ∼3-fold increase in MFF SUMO2/3-ylation (Fig 5B, D, E). Ser^155^ phosphorylation was also increased for MFF-K151R (Fig 5C), again consistent with MFF phosphorylation occurring upstream of SUMOylation. Western blots for total AMPK and phosphorylated AMPK (P-AMPK) in lysates of these CCCP experiments confirmed AMPK activation. To test whether mitophagy is being activated following CCCP, HEK293T cells were treated with CCCP (10µM and 100µM, 1hr) and blotted for LC3 (Fig S4A). No change in LC3-II (lipidated LC3, a marker of autophagosome formation) between control and CCCP at 10µM was observed, whereas at 100µM, LC3-II is increased. This confirms that under our CCCP treatment conditions, mitophagy is not being activated.

**Figure 5.**
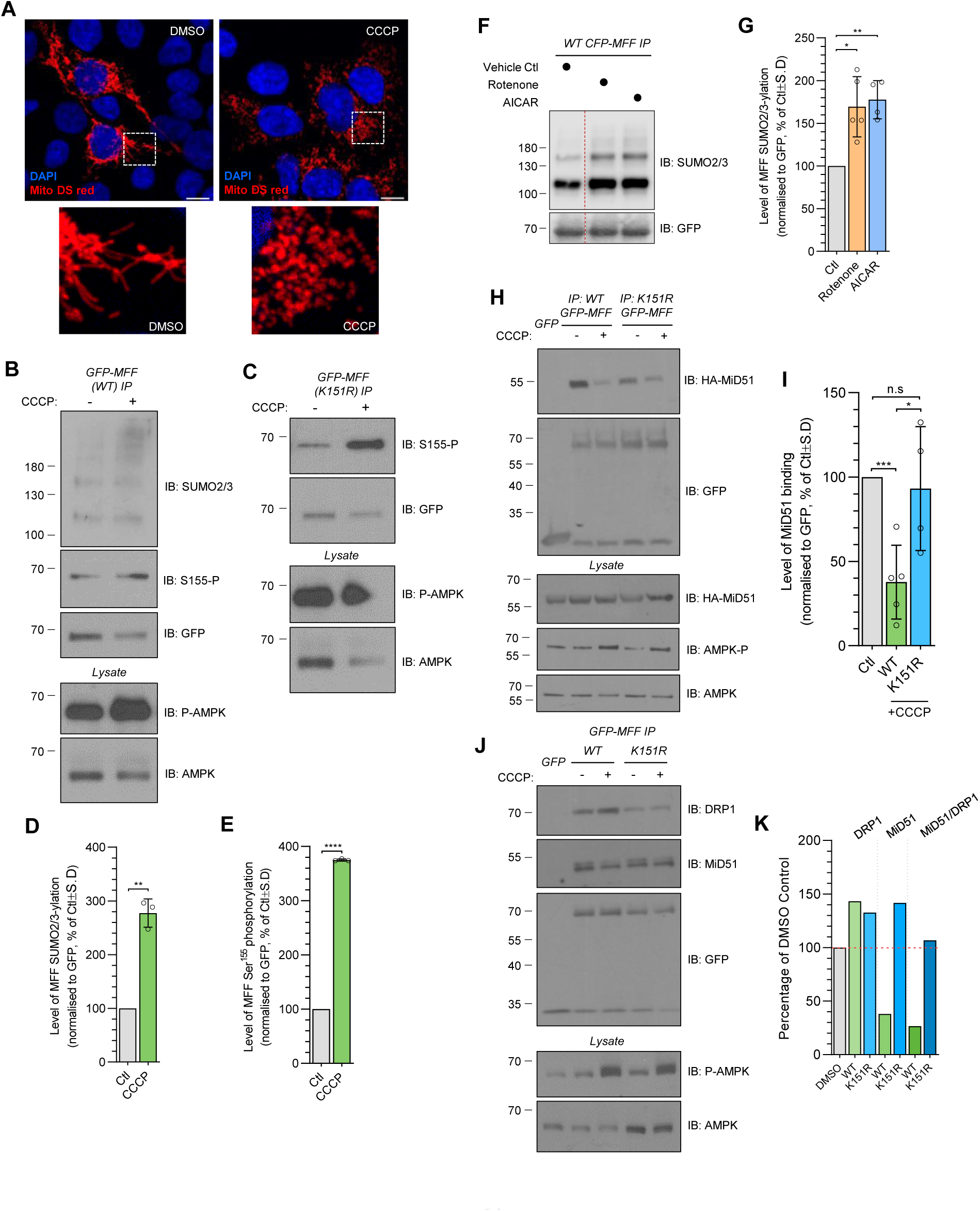
AMPK activation enhances MFF SUMOylation and MiD51 displacement in response to CCCP. **(A)** Confocal images of HEK293T cells transfected with mito DS-Red following 1hr treatment with 10µM CCCP. Scale bar 10µm. **(B, C)** HEK293T cells were transfected with GFP-MFF (WT or K151R) and treated with 10µM CCCP for 1hr before lysis. GFP-MFF immunoprecipitates were blotted for SUMO2/3, Ser^155^ phosphorylation and GFP. Input was blotted for total AMPK and AMPK activation (P-AMPK). **(D, E)** Quantification of MFF phosphorylation and SUMOylation in response to CCCP treatment, expressed as mean percentage of control ± S.D, n=3. **(F)** HEK293T cells were transfected with CFP-MFF (WT) and treated with 250ng/mL rotenone or 1mM AICAR for 1hr prior to lysis. GFP immunoprecipitates were blotted for SUMO2/3 and GFP. Quantification presented in G, expressed as mean percentage of control ± S.D, n=5 (rotenone) and n=4 (AICAR). Uncropped blot shown in Fig S4B. **(H)** HEK293T cells expressing GFP-MFF (WT or K151R) and MiD51-HA were treated with CCCP (10µM, 1hr). Co-immunoprecipitates were immunoblotted for HA and GFP. **(I)** Quantification of MiD51-HA binding to MFF following CCCP treatment, expressed as mean percentage of control ± S.D, n=4/5. **(J)** HEK293T cells expressing WT or K151R GFP-MFF were treated with 10µM CCCP for 1hr. GFP co-immunoprecipitates were probed for endogenous DRP1 and MiD51. Quantification shown in K, showing percentage of control. One-sample t-test for CCCP conditions vs vehicle controls, two-sample t-test for WT vs K151R CCCP conditions. p*<0.05, p**<0.01, p***<0.005, p****<0.0001.

To further substantiate the role of AMPK in this pathway, we next quantified MFF SUMOylation in response to treatment with the AMPK activator AICAR, and the complex I inhibitor rotenone, a specific mitochondrial inhibitor and activator of AMPK, both of which have previously been shown to lead to MFF phosphorylation (Toyama et al., 2016). Both AICAR and rotenone significantly enhanced MFF SUMO2/3 conjugation (Fig 5F, G), indicating mitochondrial stress, either via CCCP or rotenone, activates AMPK, leading to phosphorylation and subsequent SUMOylation of MFF. Moreover, our AICAR results indicate that specific activation of AMPK is sufficient to enhance MFF SUMOylation.

Because MiD51 inhibits MFF-induced activation of DRP1 GTPase activity (Osellame et al., 2016) we investigated MiD51 binding to MFF following CCCP treatment. CCCP significantly reduced the MFF-MiD association (Fig 5H, I) consistent with our finding that MFF-2SD associates less with MiD proteins (Fig 4C-F). Importantly, MiD51 binding to non-SUMOylatable MFF-K151R was not reduced by CCCP treatment. Similar results were observed for endogenous MiD51 (Fig 5J, K). Comparison of MiD51 to DRP1 in MFF-WT and MFF-K151R complexes show that the MiD51/DRP1 ratio is reduced by >70% in the MFF-WT complex following treatment with CCCP, whereas there was no change in the composition of the MFF-K151R complex (Fig 5K). Consistent with the data shown in figure 4, these results indicate that mitochondrial stress enhances MFF phosphorylation and SUMOylation, and reduces binding to MiD51 in an MFF SUMOylation-dependent manner.

### MFF SUMOylation is not required for DRP1 recruitment, but is required for CCCP-induced fragmentation

Our data support a model whereby enhanced MFF SUMOylation in response to AMPK activation promotes fission by displacing inhibitory MiD proteins from the trimeric DRP1-MiD-MFF fission complex (Fig 4H). We interrogated this model further using wild-type and MFF-KO MEF cells. Cells were treated with 10µM CCCP for 1hr to induce fragmentation and then the mitochondria imaged. In agreement with previous reports (Losón et al., 2013), wild-type MEF cells exhibit extensive fragmentation in response to CCCP, whereas MFF-KO MEF cells were resistant to CCCP-induced fragmentation (Fig S5A). We quantified the extent of fragmentation using the free-end index, which revealed a severe impairment in fragmentation in the MFF-KO cells (an increase of 79.6% vs 38.3%, wild-type vs MFF-KO, Fig S5C). We confirmed that GFP-MFF WT can induce fission when expressed in the MEF MFF-KO cells. Although not a complete rescue to wild-type MEF levels (likely due to slightly low expression levels and/or MEF cells expressing multiple isoforms of MFF (see Fig 2B)), GFP-MFF WT was sufficient to increase stress-induced fragmentation above MFF-KO levels (Fig S5B, C).

To determine the role of MFF SUMOylation in this process, we next virally expressed either GFP-MFF WT or K151R in MEF MFF-KO cells (as in Fig 2) and pretreated cells with mitotracker, treated cells with 10µM CCCP for 1hr, and stained for endogenous DRP1 (Fig 6A). Colocalisation analysis of DRP1 with mitotracker indicated that both MFF-WT and MFF-K151R recruit equivalent levels of DRP1 to mitochondria following CCCP treatment (Fig 6B). Moreover, DRP1 recruitment was not impaired in the MFF-KO cells expressing GFP alone, indicating MFF is not necessary for the recruitment of DRP1 to mitochondria following CCCP-induced stress.

**Figure 6.**
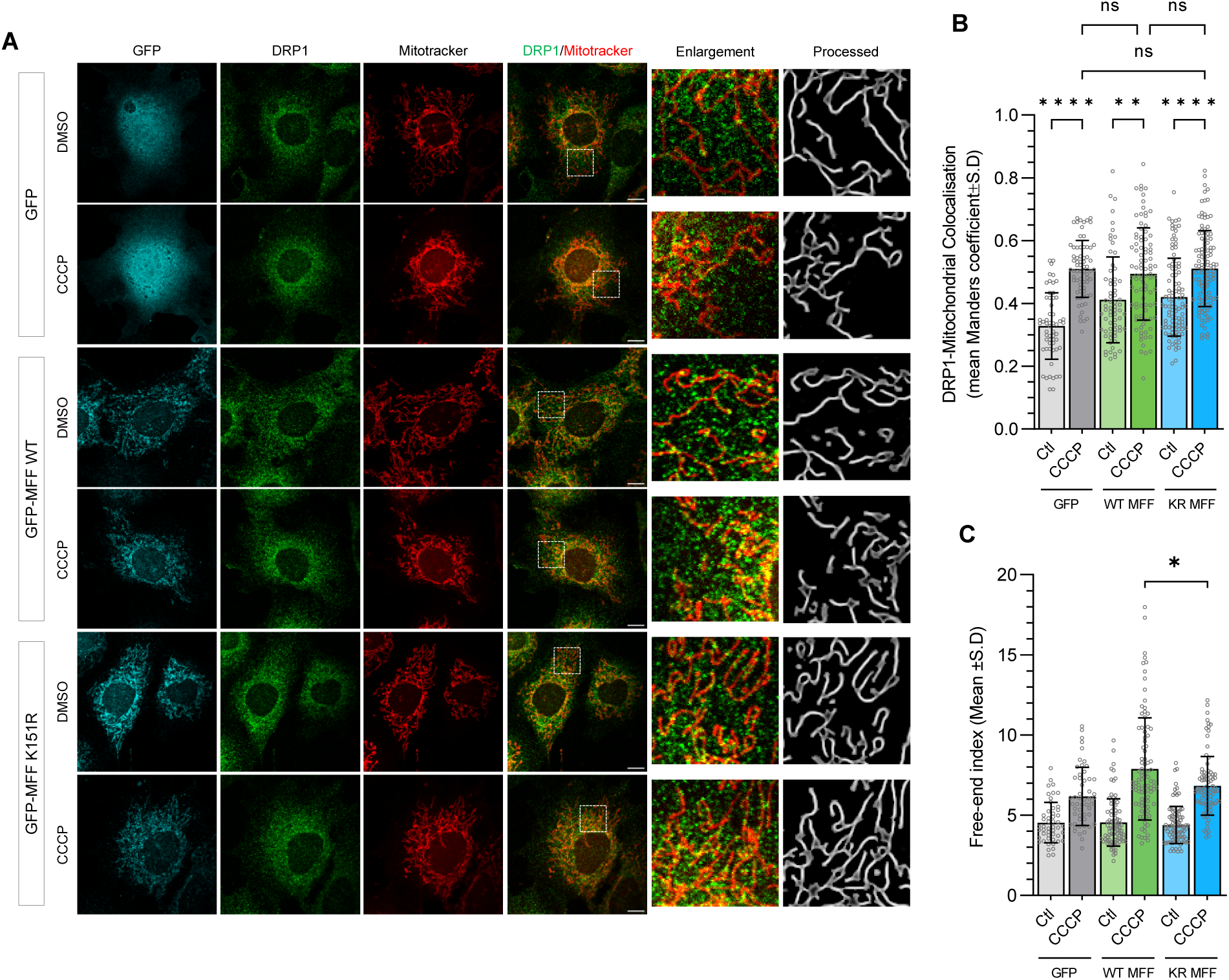
MFF SUMOylation is not necessary for DRP1 recruitment under CCCP treatment but is required for promoting mitochondrial fragmentation. **(A)** Confocal imaging of CCCP-induced mitochondrial fragmentation in MEF-KO MEF cells virally expressing GFP alone, or WT or K151R GFP-MFF. Cells were treated with CCCP (10µM, 1hr). Mitochondria stained using mitotracker deep red, endogenous DRP1 stain in green, cyan shows GFP channel. Processed images of mitochondrial stain with enlargements of highlighted area. Scale bar 10µm. **(B)** Quantification of mean Manders’ colocalisation analysis of DRP1 with mitotracker. Kruskal-Wallis test, 57-119 cells imaged from three independent experiments, p**<0.01, p****<0.0001. **(C)** Quantification of the free-end index, data generated from three independent experiments, 88-90 cells imaged (for GFP-MFF expressing cells), two independent experiments, 57-69 cells imaged for GFP expressing cells. Mann-Whitney test, p*<0.05.

Quantification of the free-end index revealed that expression of MFF-K151R impaired the fragmentation response following CCCP treatment, which was significantly lower than that observed in MFF-WT expressing cells (7.9 vs 6.8, an increase from control levels of 73.4% vs 55.8% for WT vs K151R, respectively, Fig 6C). Taken together, these data demonstrate that cells lacking MFF SUMOylation do retain the ability to recruit DRP1, and that MFF SUMOylation is an important step to promote a full stress-induced mitochondrial fission response.

## Discussion

How cells couple mitochondrial dynamics to their bioenergetic state is a fundamental question in cell biology. While AMPK-mediated phosphorylation of MFF has been shown to drive mitochondrial fission in response to bioenergetic stress (Toyama et al., 2016), the molecular events underpinning this process have remained elusive. Here, we show that MFF SUMOylation at Lys^151^ plays a key role in stress-induced mitochondrial fission. AMPK-mediated phosphorylation enhances MFF SUMOylation, which promotes mitochondrial fission by displacing the inhibitory MiD proteins from the fission complex. Our findings offer a mechanism of how mitochondrial fission complexes fine-tune the relative ratios of MFF to MiD to dynamically regulate fission under differing conditions.

More specifically, we assessed DRP1 recruitment and mitochondrial morphology following re-expression of MFF-WT or MFF-K151R in MFF-KO cells. Our results indicate that MFF SUMOylation is not required for DRP1 engagement with the mitochondria or to regulate mitochondrial morphology under basal conditions. These findings are consistent with compensation by other DRP1 receptors that can independently recruit DRP1 and promote fission (Palmer et al., 2013; Osellame et al., 2016; Losón et al., 2013). As previously reported, MFF-KO cells exhibit resistance to CCCP induced mitochondrial fragmentation (Losón et al., 2013; Otera et al., 2010; Osellame et al., 2016), confirming that MFF is a core component of the fission machinery. However, DRP1 recruitment increases under CCCP conditions in the MFF-WT and MFF-K151R expressing cells (Fig 6B) indicating that MFF SUMOylation is not required for DRP1 recruitment under stress. Importantly, while WT-MFF re-expression was able to rescue the fragmentation phenotype, fragmentation was still impaired in MFF-K151R expressing cells. This demonstrates that when MFF cannot be SUMOylated fission is impeded downstream of DRP1 engagement.

Mitochondrial morphology is regulated by balanced fusion and fission (Detmer and Chan, 2007; Twig et al., 2008). Therefore, a fragmented mitochondrial network may be a result of enhanced fission, or alternatively, reduced fusion. MFF is well established as a pro-fission protein, mediating DRP1-dependent fission (Losón et al., 2013; Otera et al., 2010), which does not negatively regulate fusion (Gandre-Babbe and van der Bliek, 2008). Thus, our data strongly support a direct role of MFF SUMOylation in mitochondrial fission, and not inhibition of fusion.

To examine the role of MFF SUMOylation in stress-induced fission, we used AMPK phosphorylation mutants of MFF (2SA and 2SD). Our results demonstrate that MFF phosphorylation does not enhance DRP1 association with MFF *per se*, as has been previously postulated (Toyama et al., 2016), but rather MFF SUMOylation controls the stoichiometry of MiD proteins within the trimeric DRP1-MiD-MFF fission complex. The high phospho/SUMO state (MFF-2SD) binds less to MiD proteins, whereas the non-SUMOylatable mutant (MFF-K151R) and low phospho/SUMO state (MFF-2SA) have enhanced binding to MiD proteins. We further confirm this model using CCCP, which enhances MFF SUMOylation and reduces MiD51 binding, whereas MiD51 binding to non-SUMOylatable MFF-K151R is unchanged. Importantly, Ser^155^ phosphorylation is increased following stress in both WT and K151R MFF. Thus, our data demonstrate that SUMOylation is downstream of phosphorylation, and it is the SUMOylation event at K151 that promotes MiD displacement from the fission complex.

We next used protein crosslinking approaches and immunoprecipitation of MFF mutants to show that the DRP1-MiD-MFF fission complexes assemble with DRP1 in a range of oligomeric states (monomer, tetrameric, higher order), regardless of the post-translational modifications of MFF. These data contrast, in part, with the findings of Yu and colleagues, who reported that MiD proteins can associate with a wide range of active and inactive states of DRP1, whereas MFF favours active forms and higher order states (Yu et al., 2021). This apparent discrepancy might be due to differences in the time period used for DSS crosslinking or other experimental details such as the conditions used to isolate the fission complex. In particular, under our conditions, we isolate the trimeric complex containing MiD, which likely explains our detection of a wide range of DRP1 states. Nonetheless, the findings by Yu and colleagues align with the suggestion that the MiD proteins act as a scaffold for inactive forms of DRP1, as previously suggested (Palmer et al., 2011), whereby they recruit DRP1 from the cytosol, and following a fission stimulus, transfer DRP1 to MFF, forming a fission competent complex (Yu et al., 2021). Our data significantly refines this model to explain how post-translational modifications of MFF regulate the relative levels of MiD to DRP1 to promote fission under stress (working model in Fig 7). Exactly how the complex is rearranged and DRP1 transferred to MFF remains an important unanswered question.

**Figure 7.**
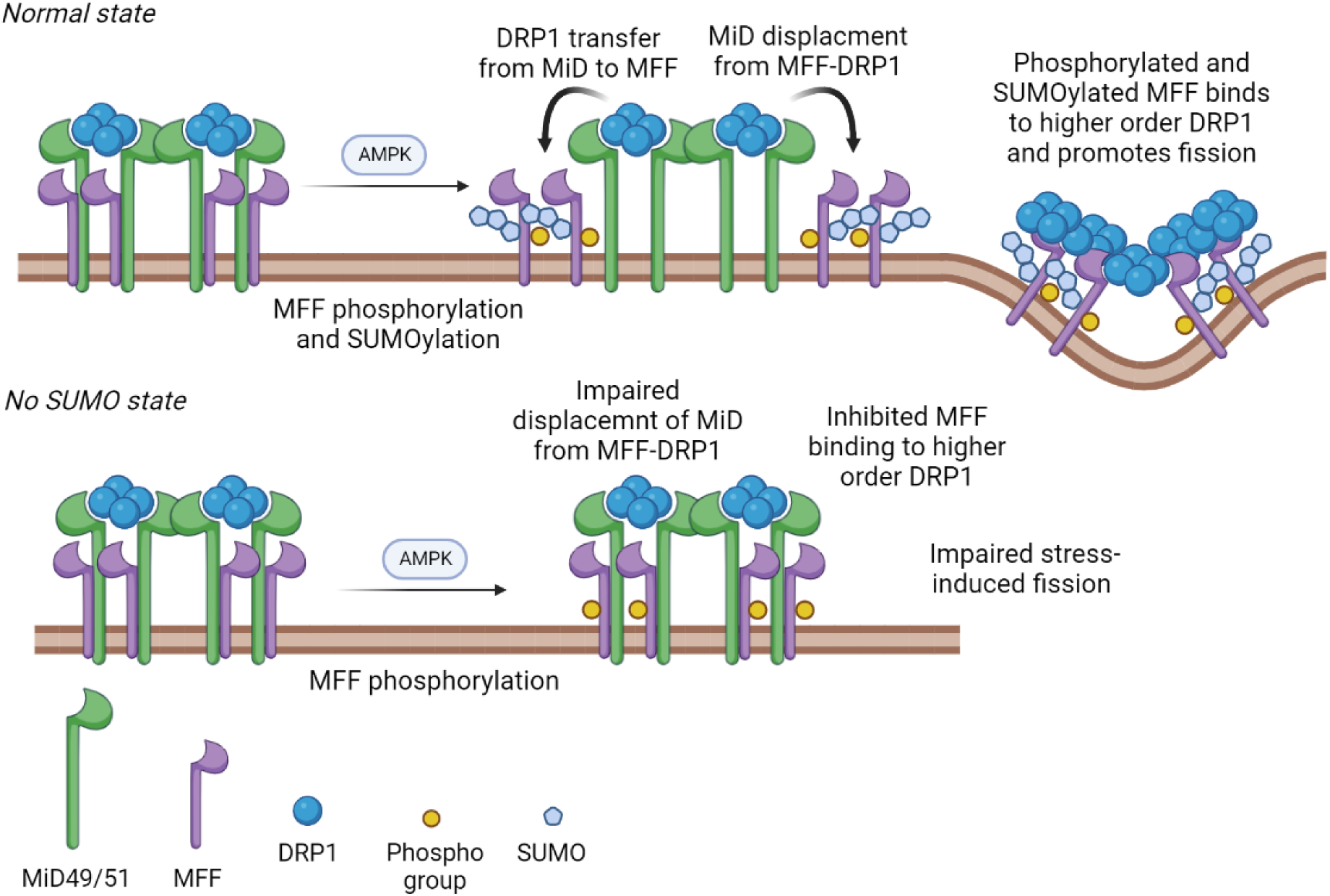
Working model of MFF-SUMOylation dependent stress-induced fission. MFF, DRP1 and MiD49/51 proteins exist in a trimeric complex. Upon AMPK activation, MFF is phosphorylated at Ser^155^ and Ser^172^, leading to MFF SUMOylation at Lys^151^ (for simplicity, only one phosphorylation site is shown). In the wild-type condition, this results in reduced MiD association and displacement from the complex, leading to the formation of MFF-DRP1 fission competent complexes. How MiD proteins transfer DRP1 to MFF remains to determined. When MFF cannot be SUMOylated, DRP1 is still recruited, and MFF is still phosphorylated, but MiD proteins remain associated in the trimeric complex. MFF-DRP1 fission complexes are not efficiently formed, leading to impaired fragmentation in response to stress. Created with BioRender.com.

In addition to AMPK phosphorylation promoting MFF SUMOylation, we identify MAPL as a SUMO E3 ligase of MFF, and SENP3 and SENP5 as MFF deSUMOylating enzymes (Fig S2F, G). MAPL mediates DRP1 SUMOylation with SUMO1, which is important for DRP1 stability and driving fission (Braschi et al., 2009; Harder et al., 2004; Zunino et al., 2007) as well as facilitating mitochondrial-endoplasmic reticulum contact during apoptosis (Prudent et al., 2015). Overexpression of SENP5 results in tubulated mitochondria, reduced DRP1 levels and diminished SUMO conjugation in mitochondrial fractions (Zunino et al., 2007). These results support a hypothesis that general mitochondrial protein SUMOylation favours fission, which is consistent with our observations that MFF SUMOylation promotes fission.

On the other hand, DRP1 is also SUMOylated by SUMO2/3, which sequesters DRP1 in the cytosol and impairs the DRP1-MFF interaction during oxygen-glucose deprivation, a modification reversed by SENP3 (Guo et al., 2017, 2013). Taken together, our findings and these reports highlight central roles of mitochondrial protein SUMOylation under basal and stress conditions. The interrelationship between SUMOylation of DRP1 and MFF (by SUMO1 and/or SUMO2/3), and how MAPL and SENPs regulate their SUMOylation status to dynamically control appropriate mitochondrial responses, represent interesting and important avenues for future work.

Intriguingly, there is evidence that post-translational modifications of mitochondrial proteins play key roles in regulating mitochondrial fission during cell division. During mitosis it has been reported that protein kinase D (PKD) phosphorylates MFF at Ser^155^, Ser^172^ and Ser^275^, modifications necessary and sufficient for mitochondrial fission and correct chromosome segregation during cell division (Pangou et al., 2021). Importantly, this process is independent of AMPK phosphorylation, indicating that MFF has at least two different kinases that act on the same sites during distinct cellular processes.

These findings raise the interesting question of whether PKD-MFF and AMPK-MFF pathways promote mitochondrial fission under different cellular conditions through SUMOylation of MFF. For example, does PKD-mediated phosphorylation lead to enhanced MFF SUMOylation during cell division? Conversely, SENP5 is primarily a nuclear enzyme that translocates to mitochondria during mitosis where it deSUMOylates DRP1 to promote fission during cell division (Zunino et al., 2009). It is unknown if SENP5 deSUMOylates MFF during cell division.

We show that MFF is modified by both SUMO1 and SUMO2/3, exists in multiple SUMO states and exhibits multiple lengths of SUMO chains. Another poly-SUMO substate is PML (Tatham et al., 2001), an important component of nuclear bodies; multi-protein assemblies within the nucleus which regulate a number of key nuclear functions, such as transcription, DNA repair and senescence (Lallemand-Breitenbach and de Thé, 2010). PML SUMOylation is essential for correct nuclear body localisation and formation (Fu et al., 2005; Zhong et al., 2000). Distinct SUMO-interacting proteins are recruited to the poly-SUMO chain on PML (Erker et al., 2013; Lallemand-Breitenbach et al., 2008) and exhibit a binding preference depending on the SUMO chain composition (Sriramachandran et al., 2019). This raises the question of whether the multiple SUMOylation states of MFF, and the different chain compositions, could potentially act to recruit distinct proteins that recognise the SUMO chain and perform different functions. Indeed, other proteins involved in fission/fusion have functions beyond their primary role; Fis1 binds to Bap31 on the endoplasmic reticulum, with roles in apoptosis (Iwasawa et al., 2011), and Mfn2 tethers the mitochondria and endoplasmic reticulum to form stable inter-organelle contact sites and mediate calcium transfer (de Brito and Scorrano, 2008). Future research into the composition and potential interactors of the MFF poly-SUMO chain will yield greater understanding of the role of poly-SUMO chains and potential functions beyond fission.

Fission complexes incorporating MFF have been reported to play a major role in mitochondrial biogenesis and cell proliferation, whereas those containing Fis1 are involved in mitochondrial fission in response to mitochondrial damage (Kleele et al., 2021). While such demarcated roles for MFF and Fis1 receptors is intriguing, it is not consistent with observations that MFF silencing protects against stress-induced fragmentation (Osellame et al., 2016; Otera et al., 2010; Losón et al., 2013) nor with our data indicating a key role for MFF-SUMOylation in stress-induced fission. One explanation could be that the report by Kleele and colleagues used UV irradiation as a stressor, while our work primarily used CCCP. Thus, it is possible that different pathways and fission machinery are activated under different conditions. Additionally, mitochondrially targeted AMPK (mito-AMPK), which differs in its spatial activation across the mitochondrial network in response to energetic stress, suggests a highly regulated system of AMPK responses to local microenvironments at the mitochondrial surface (Drake et al., 2021). Together, these findings suggest a picture of the local and precise regulation of fission in response to local energy demands. Mito-AMPK is activated in defined regions within the mitochondrial network that are exposed to higher AMP/ATP ratios. The resultant spatially restricted phosphorylation and SUMOylation of MFF causes MiD protein displacement from, and the relief of inhibition of, local fission complexes. In this way, spatially regulated changes within the network allow mitochondria to make appropriate fission responses to localised stressors.

Thus, we propose that MFF SUMOylation is tightly controlled, and likely exists as a continuum, rather than an all-or-nothing response. In this way the SUMOylation status of MFF can act to fine-tune and couple mitochondrial fission with the bioenergetic state of the cell. Future investigation into the conditions and the pathways involved in regulating the SUMOylation state of MFF will provide insight into the nuanced regulation of the fission machinery.

In conclusion, we show that MFF SUMOylation is a critical step in stress-induced mitochondrial fission. We propose a mechanistic model of fission in which MFF SUMOylation modulates the stochiometric composition of the DRP1-MiD-MFF trimeric fission complex to dynamically regulate the fusion/fission balance to rapidly induce fragmentation during bioenergetic stress.

### Materials and methods Reagents and antibodies

Dulbecco’s Modified Eagle’s Medium (DMEM, Lonza), heat-inactivated foetal bovine serum (FBS, Sigma), penicillin and streptomycin (Gibco), 0.05% trypsin-EDTA (Gibco). Lipofectamine 2000 was from ThermoFisher. Poly-L-lysine (PLL), rotenone (dissolved in DMSO), carbonyl cyanide *m*-chlorophenyl hydrazone ((CCCP), dissolved in DMSO), β-glycerophosphate, Na-pyrophosphate, N-ethylmaleimide (NEM), EDTA, triton-X and glycerol were from Sigma. AICAR (dissolved in cell culture grade H_2_O) was from Tocris. DSS was from ThermoFisher, prepared fresh in DMSO. Protease inhibitor (complete, EDTA-free, protease inhibitor cocktail tablets) were from Roche. Glutathione sepharose beads were from GE Healthcare Life Sciences, GFP-Trap beads were from Chromotek, anti-SUMO2/3 beads were from Cytoskeleton.

For Western blotting, HRP-conjugated anti-mouse (raised in goat), anti-goat (raised in rabbit), anti-rabbit (raised in goat) and anti-rat (raised in rabbit) were obtained from Sigma and used at a dilution of 1:10,000. Primary antibodies for Western blotting are listed in table 1. Primary antibodies used for imaging were mouse anti-DRP1 (BD Bioscience, #611113, at 1:400) and chicken anti-GFP (Abcam, #13970, at 1:1000). Secondary antibodies for imaging were Cy2 anti-chicken and Cy3 anti-mouse (raised in donkey) from Jackson ImmunoResearch and used at 1:400. Mitotracker deep red was obtained from Invitrogen (#M22426), diluted in DMSO and used at a final concentration of 100nM.

**Table 1.** List of antibodies, supplier and catalogue numbers used in this study

| **Protein** | **Supplier** | **Reference number** | **Species** |
| --- | --- | --- | --- |
| AMPK α-1 | ThermoFisher | AHO1332 | Mouse |
| P-AMPK (T172) | Cell Signalling | 2535S | Rabbit |
| p-(S/T) AMPK substrate motif | Cell Signalling | 5759S | Rabbit |
| DRP1 | BD BioScience | 611113 | Mouse |
| FLAG | Sigma | F3165 | Mouse |
| GFP | Chromotek | pabg-1-100 | Rat |
| GST | GE Healthcare | 27457701V | Goat |
| HA | Sigma | H3663 | Mouse |
| LC3A/B | Cell Signalling | 4108 | Rabbit |
| MiD49 | Proteintech | 16413-1-AP | Rabbit |
| MiD51 | Proteintech | 20164-1-AP | Rabbit |
| MFF | Santa Cruz | SC-398731 | Mouse |
| SENP3 | Cell Signalling | 5591S | Rabbit |
| SENP5 | Abcam | ab58420 | Rabbit |
| MAPL | Abcam | ab155511 | Rabbit |
| SUMO1 | Cell Signalling | 4930S | Rabbit |
| SUMO2/3 | Cell Signalling | 4971S | Rabbit |
| Ubiquitin | Cell Signalling | 3936 | Mouse |
| β-actin | Sigma | A5441 | Rabbit |

### Plasmids and siRNA

A pool of siRNA against MAPL (Mul1) was purchased from Dharmacon™ (ON-TARGETplus human MUL1 siRNA). Control pool of siRNA was from Dharmacon™ (ON-TARGETplus Non-targeting Pool). Both were dissolved in RNAse-free water and used at a final concentration of 20nM. Human SENP siRNAs were from Sigma (SENP3: ACGUGGACAUCUUCAAUAA, SENP5: AAGUCCACUGGUCUCUCAUUA, Control targeting luciferase: CUUACGCUGAGUACUUCGA) and used at 100nM. GST-tagged DRP1 receptors have been described previously (Guo et al., 2017). CFP-MFF was constructed by subcloning the MFF human isoform 1 sequence from GST-MFF into the BamHI/HindIII sites of pECFP-C1. MitoDS Red (pDsRed2-mito) was from Clontech.

3xFLAG-SUMO1 and SUMO2 were produced by PCR-based cloning of human SUMO1 or SUMO2 into the BamHI site of one of the pCMV-Flag series of vectors, using the primers hSUMO1 BamHI F (CTCGGATCCATGTCTGACCAGGAGGCAAAA) and hSUMO1 BamHI R (CACGGATCCTAACCCCCCGTTTGTTCCTG) or hSUMO2 BamHI F (CTCGGATCCATGGCCGACGAAAAGCCCAAG) and hSUMO2 BamHI R (CACGGATCCTAACCTCCCGTCTGCTGTTG). GFP-Fis1 was produced by PCR-based cloning of rat Fis1 (amplified from p3xFLAG-CMV-10-Fis1, a kind gift from Michael Schrader (University of Exeter, UK)) into the HindIII and EcoRI sites of pEGFP-C3 (Clontech), using the primers rFis1 HindIII F (CTCAAGCTTATGGAAGCCGTGCTGAACGAG) and rFis1 EcoR1 R (GTGGAATTCCCTTCAGGATTTGGACTTGGACAC). Mutants of MFF were generated by KOD Hot Start DNA Polymerase PCR-based site-directed mutagenesis (see table 2 for primers).

**Table 2.**
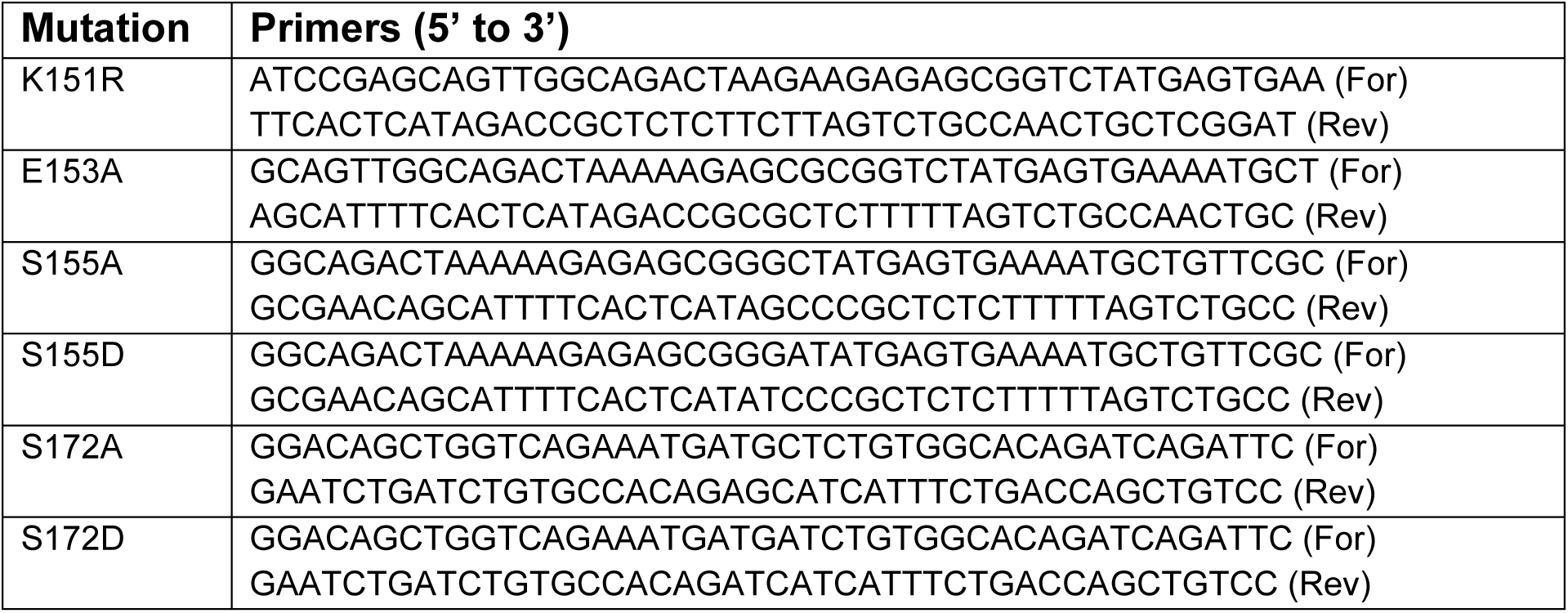
Primers for site-directed mutagenesis

Lentiviral GFP-MFF constructs were produced in the plasmid pXLG3-PX-GFP-WPRE. pAcGFP-tagged human MFF isoform 1 was subcloned from pAcGFP-C1-MFF (a kind gift from Gia Voeltz (Addgene plasmid #49153) by digestion of pAcGFP-C1-MFF with NheI and BamHI to isolate the pAcGFP-MFF insert, and ligating into SpeI and BamHI cut pXLG3-PX-GFP-WPRE, in place of the GFP. The MFF K151R mutant was produced in exactly the same way after first mutating K151 to arginine in pAc-GFP-C1-MFF, as described below.

### Generation of MiD-HA and MiD-GFP

RNA was extracted from HEK293T cells using Qiagen RNeasy mini kit as per the manufacturer’s instructions. Briefly, a confluent 6cm dish of HEK293T cells was scraped into 600µL RLT buffer supplemented with β-mercaptoethanol and centrifuged at 16,000rcf. Supernatant was precipitated using an equal volume of 70% ethanol and transferred to a RNeasy Mini Spin column. Column was washed and RNA eluted into an RNAse free microcentrifuge tube with 25µL RNase-free water. cDNA was synthesised from RNA using RevertAid First Strand kit (ThermoFisher) according to manufacturer’s protocol.

The primers in table 3 were used to amplify the complete coding sequence of MiD49 and MiD51 from cDNA and subcloned into the BamH1/HindIII sites of pEGFP-N1 and pcDNA3.1 (for MiD-GFP and MiD-HA, respectively). The fidelity of all constructs was confirmed by DNA sequencing (Eurofins Genomics).

**Table 3.**
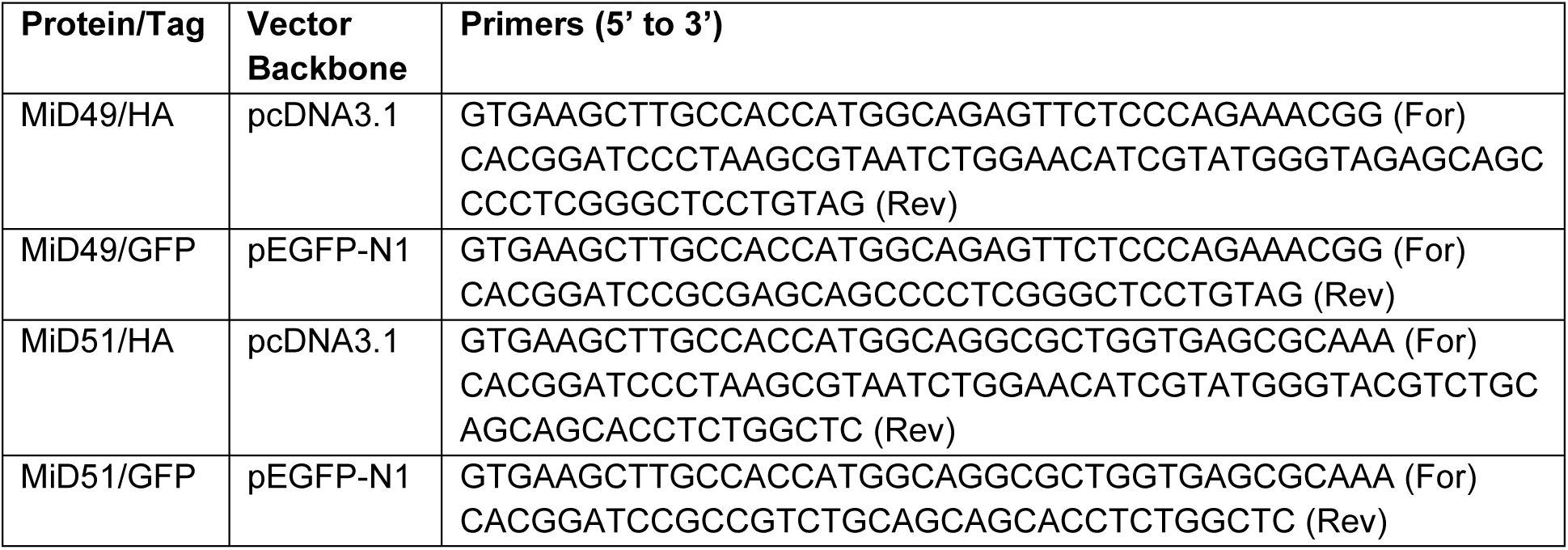
Primers for generation of MiD49/51 constructs

### Cell culture and transfection

Human Embryonic Kidney cells (HEK293T) cells were from the European Collection of Cell Cultures (ECACC). Mouse Embryonic Fibroblasts (MEF, WT and MFF-KO) have been described previously (Losón et al., 2013). HEK293T and MEF cells were cultured in DMEM containing 4mM L-glutamine, supplemented with 10% FBS, streptomycin (100µg/mL) and penicillin (100 units/mL). Cells were maintained in a humidified atmosphere of 5% CO_2_ at 37°C. For transfection experiments, 6cm dishes were coated with 0.1mg/mL PLL. HEK293T cells were seeded at 1.5×10^6^ cells per dish in 4mL transfection media (culture media lacking antibiotics). The following day, HEK293T cells were transfected using Lipofectamine 2000 and incubated for 36-48hrs. siRNA was transfected along with plasmid DNA using Lipofectamine as above.

### Lentivirus production and transduction

Lentiviral particles were produced in HEK293T cells. Briefly, 2×10^6^ cells were seeded into a 6cm dish. The next day, cells were transfected with 4µg pXLG3-based lentiviral plasmid, 3µg and 1µg of the helper plasmids p8.91 and pMD2.G, respectively, using polyethylenimine (PEI; Sigma). Transfection mixtures were left on for 4 hours, before replacement with 3mL complete DMEM. Culture media, containing lentiviral particles, was collected 48h later, centrifuged at 2500rcf to pellet cell debris, and filtered through a 0.45µm syringe filter. Virus-containing supernatant was then aliquoted into 500µL aliquots and frozen at -80°C. For cell transduction, lentivirus was thawed, and the desired amount added drop-wisely to the cells being transduced. Transduced cells were passaged several times and used in experiments as appropriate.

### SDS-PAGE and Immunoblotting

10-12% polyacrylamide gels were made in house and samples resolved by SDS-PAGE. Proteins were transferred to PVDF membranes (Millipore), blocked in 5% milk or 4% BSA (prepared in PBS-T) for 1hr at room temperature and incubated with primary antibody for either 1hr at room temperature or overnight at 4°C (table 1 for primary antibodies). Membranes were washed with PBS-T and incubated with secondary antibody conjugated to horseradish peroxidase at 1:10,000. Membranes were washed in PBS-T and assayed for chemiluminescence by using ECL and X-ray film (ThermoFisher) or using a Li-COR Odyssey Fc scanner (Fig 5F and S3D).

### Immunoprecipitations and GST pulldowns

For immunoprecipitation experiments, cells were washed in ice cold PBS and lysed on ice for 45-60 minutes in the following lysis buffer: 20mM tris (pH 7.4), 137mM NaCl, 1% triton X-100, 10% glycerol, 25mM β-glycerophosphate, 2mM Na-pyrophosphate, 20mM NEM, 2mM EDTA, supplemented with protease inhibitors. For investigations into covalent modification (i.e. Ser^155^ phosphorylation, SUMO conjugation), lysis buffer was supplemented with 0.1% SDS and samples briefly sonicated. Lysate was clarified for 20 minutes at 16,000rcf at 4°C. Supernatant was collected and kept on ice. 4% input was taken and 1 volume of 2× Laemmli sample buffer added before heating at 95°C for 10 minutes. On the remaining lysate, GFP-TRAP beads (Chromotek) were used to perform immunoprecipitations of GFP and CFP-tagged proteins and glutathione-sepharose 4B beads (GE Healthcare Life Sciences) were used for pulldowns of GST-tagged proteins. Lysate was added to washed beads and incubated at 4°C for 90 minutes with slow rotation. Beads were washed three times (in lysis buffer without proteases inhibitors, SDS or NEM). After final wash 2x Laemmli buffer was added, and samples were boiled at 95°C for 10 minutes.

For enrichment of SUMOylated proteins anti-SUMO2/3 antibody conjugated beads (Cytoskeleton), or control beads protein G beads (Cytvia) were used according to manufacturer’s protocol. Briefly, HEK293T cells were lysed in lysis buffer as above, supplemented with 4% SDS and 20mM NEM (or equivalent amount of H_2_O for control conditions). Lysis initially performed at room temperature for 5 minutes, lysate diluted to 0.1% SDS and then placed on ice for 30 minutes. 0.5mg (0.5mg/mL) of lysate was incubated with 30µL beads overnight at 4°C. Beads were washed three times and then boiled in 2x Laemmli buffer.

For chemical cross-linking prior to co-IP experiments, DSS was used as previously described (Yu et al., 2021), with slight modification. Briefly, transfected HEK293T cells were washed in PBS (containing 1mM CaCl_2_ and 0.5mM MgCl_2_) and incubated with 1mM DSS at room temperature for 30 minutes, and then quenched in 50mM Tris (pH 7.5) for 15 minutes at room temperature. Cells were washed in PBS, lysed in 50mM Tris (pH 7.5), 150mM NaCl, 1% triton, 20mM NEM, supplemented with protease inhibitors. Lysate was clarified, and GFP-IP performed on supernatant as described above. Samples were resolved by SDS-PAGE using a pre-cask 4-20% gradient gel (Bio-Rad).

### *In vitro* deSUMOylation and deubiquitination assay

The catalytically active domain of SENP1 (produced as described previously (Rocca et al., 2017)) and purified USP2 (a kind gift from the Ron Hay lab (University of Dundee, UK)) was used to enzymatically remove SUMO and ubiquitin from MFF, respectively. To obtain sufficient material, multiple 6cm dishes of HEK293T cells transfected with WT CFP-MFF or CFP (negative control) were washed in PBS and pooled together in lysis buffer containing 0.1% SDS, protease inhibitors and 20mM NEM on ice. CFP-MFF was then immunoprecipitated on GFP-TRAP beads as per immunoprecipitation protocol. Beads were washed three times in wash buffer (137mM NaCl, 50mM tris (pH 7.4), 5mM MgCl_2_) and separated equally into three fresh Eppendorf tubes, and a final concentration of 100nM GST-SENP1 or 500nM GST-USP2 was added for 2hrs at 37°C, with occasional agitation (wash buffer added to control and CFP conditions). An equal volume of 2x Laemmli buffer was then added and samples were boiled at 95°C for 10 minutes.

### Total cell lysis

For cell lysis in Fig 1B, transfected HEK293T cells were washed in 1x PBS and lysed in the following buffer: 50mM tris (pH7.4), 137mM NaCl, 1% triton, protease inhibitors, and either 2% SDS, or equivalent volume of H_2_O. Samples were lysed (initially at RT for 5 minutes, then kept on ice for 30 minutes), sonicated, an equal volume of 2x Laemmli buffer added, and boiled at 95°C for 10 minutes. For cell lysis, as in Fig 2B and 2H, MEF cells were grown in 6 well plates, washed in 1x PBS, lysed in 1x Laemmli buffer and boiled at 95°C for 10 minutes.

### Densitometry analysis of Western blots

• X-ray films were scanned as PNG files and analysed using ImageJ software. Files were converted to 8-bit format, analysed using the Gel Analyser tool and area under the curve values extracted. For immunoprecipitation experiments, all values are normalised to the respective GST or GFP reprobe for unmodified tagged-MFF and expressed as a percentage of the control. Corresponding to Figs 3A-D and S2A-D: for investigating SUMO conjugation in CFP-MFF IPs, the mono-SUMO band saturated before the higher molecular weight bands were detected. Therefore, we took different exposures and performed independent analysis of the mono-SUMO and the higher molecular weight bands of SUMOylated MFF. In Fig 3C-D, the SUMO values are mean averages of the mono-SUMO and higher molecular weight species. In Figs 5F, S3D and S4, blots were developed using a Li-COR Odyssey Fc, and quantified using Li-COR Image Studio software. Cropped blots are indicated with a red dotted line (Figs 3A, 5F).

### Immunocytochemistry and Imaging

MEF cells were grown on PLL coated glass cover slips and were pre-treated with mitotracker deep red at a final concentration of 100nM for 45 minutes prior to fixation. For experiments of CCCP treatment of 1hr, cells were incubated in mitotracker dye for 45 minutes prior to treatment with CCCP. Cells were fixed in 4% formaldehyde for 15 minutes. Cells were washed three times in PBS, permeabilised with 0.1% triton X-100 (in PBS) for 3-4 minutes, then washed in PBS. Cells were incubated for 3-4 minutes in 100mM glycine/PBS to quench unreacted formaldehyde and washed once in PBS. To block non-specific binding, cells were incubated in 3% BSA/PBS for 20 minutes at room temperature. Primary antibody (DRP1 1:400, GFP 1:1000) was prepared in 3% BSA and coverslips incubated with primary antibody for 60 minutes at room temperature. Coverslips were washed three times with PBS and then incubated with secondary antibody (Anti-mouse Cy3 and Anti-chicken Cy2, prepared in 3% BSA/PBS at 1:400) for 45 minutes. Coverslips were washed four times in PBS and mounted on glass microscope slides using Fluoromount-G (containing DAPI).

Imaging was carried out using a Leica SP5-II confocal laser scanning microscope attached to a Leica DMI 6000 inverted epifluorescence microscope. Images were captured using a 63x HCX PL APO CS oil-immersion objective, with 512×512 pixel resolution and optical zoom of 3x at 400Hz. Z-stacks were taken with 0.25µm incremental steps. DAPI was excited using a 50mW 405nm diode laser, Cy2 was excited using a 150mW Ar laser (488nm), Cy3 using a 20mW solid state yellow laser (561nm) and mitotracker deep-red using a 20mW Red He/Ne (633nm). All the parameters were kept constant for a complete set of experiments.

### DRP1 colocalisation analysis

We generated a cytoplasmic mask to designate the area for DRP1-mitotracker colocalisation analysis, which would remove the nuclei and non-cytoplasmic regions, and avoid manually designating the cytoplasmic region, making analysis more objective. Using the DRP1 stain channel the following workflow was used: 1) Segmentation of the nuclei using the following steps: median filter of radius 2-pixels. Applied global threshold to binarise the image (Otsu method), removed holes in the binarised image, identified nuclei as connected regions to have a 2D area larger than 50µm^2^, using the MorphoLibJ library. 2) To identify the extent of the cell: median filter applied with radius 2-pixels. Applied global intensity threshold to binarise image (Huang method), combined the binarised nuclear and cytoplasmic images to get a complete binary image of the cell. Applied another median filter (2-pixel radius). Applied fill holes as before and used distance-based watershed transform to split any adjacent cells. Cells had a minimum 2D area larger than 400µm^2^ to be identified. 3) Subtracted the nucleus from the whole cell to yield the cytoplasmic region. Cytoplasmic objects identified as before, using connected foreground labelled pixels. Cytoplasm detected area must be larger than 50µm^2^. The 2 channels were normalised to the full 8-bit intensity range. Manders’ colocalisation was calculated using ImageJ’s coloc2 tool between the DRP1 channel and mitotracker channel within the cytoplasmic mask.

Plugin is archived in S. Cross, (November 28, 2022) “Colocalisation analysis” (Version 0.7.22) Zenodo https://zenodo.org/record/7372802#.Y4TF5hTP1D8.

### Mitochondrial morphology analysis

In order to analyse the mitochondrial network of MEF cells in an objective and quantifiable manner, we adapted a method developed by Valente and colleagues, who described a macro in conjunction with ImageJ software (Valente et al., 2017). We adjusted the pre-processing steps to better reproduce the mitochondrial morphology of our imaging and manually extracted our own parameters for analysis. Firstly, using the freehand selection tool, we outlined the cell of interest and cleared the outside. The nuclear region was also traced and excluded from analysis. The confocal z-stack was projected to a single image (max intensity), local contrast enhanced (blocksize=125 histogram bins=256 maximum slope=2) and background subtracted (radius 10, with sliding paraboloid). Following this, two filters were applied: median filter (radius 1 pixel), unsharp filter (0.4 sigma radius, 0.7 mask weight). We incorporated a plug-in called Tubeness (sigma 0.2), which we found increased the detection of smaller and dimmer mitochondria, and also prevented over fragmentation of the network, making the skeleton a more faithful representation of the raw image. Following pre-processing, the image was binarised and skeletonised (ImageJ’s make binary and skeletonise). This generates a 1-pixel outline which allocates three types of pixel, based on the immediate neighbours: end pixels have either 1 or 0 neighbours, slab pixels have 2 neighbours, whereas junctions have 3 or 4 neighbours. We used the ImageJ’s in-built analyse skeleton function, which generates a table of information on the branches and pixels. From the table of branch information, the values were exported to excel and various parameters were manually extracted: 1) Mean number of branches per network: mean number of branches within structures containing ≥2 branches. The number of branches column was arranged in numerical order, branches of less than 2 removed, and the average mean calculated. 2) Mean mitochondrial length: extracted from the average branch length, which is the length between two endpoints, an end-point and junction, or two junctions. The branch length column was arranged in numerical order, non-zero lengths were removed (i.e. single pixels), and the average mean calculated. 3) Free-end index: number of free ends as a percentage of the total number of pixels detected (sum of free ends / sum of all pixels (free ends, junctions, and slab pixels)) x 100. We used this parameter as a measure of the extent of fragmentation.

Plugins for the mitochondrial morphology analysis are archived in S. Cross, (November 28, 2022) “Preprocess” and “Branch analysis” analysis” Zenodo https://zenodo.org/record/7372802#.Y4TF5hTP1D8.

### Statistical analysis

Statistical analysis was performed using GraphPad Prism software version 8. For quantification of densitometry of Western blots, all values are presented as mean ±S.D, expressed as a percentage of control. One sample t-test was used to determine significance between conditions and control (set to 100), unpaired t-test was used to determine significance between two groups. For multiple comparisons, one-way ANOVA was used followed by Tukey’s *post hoc* test. For analysis of imaging, data was tested for normality distribution using the D’Agostino & Pearson test. If this test was passed, then a parametric test (t-test, one-way ANOVA followed by Tukey’s *post hoc* test) was used to determine significance. If failed, then a non-parametric test (Mann-Witney test, Kruskal-Wallis test followed by Dunn’s *post hoc* test) was used to determine significance. A p-value of <0.05 was considered significant. P values, independent repeats and statistical approach are described in the figure legends.

## Figures

## Supplementary data

**Figure S1.**
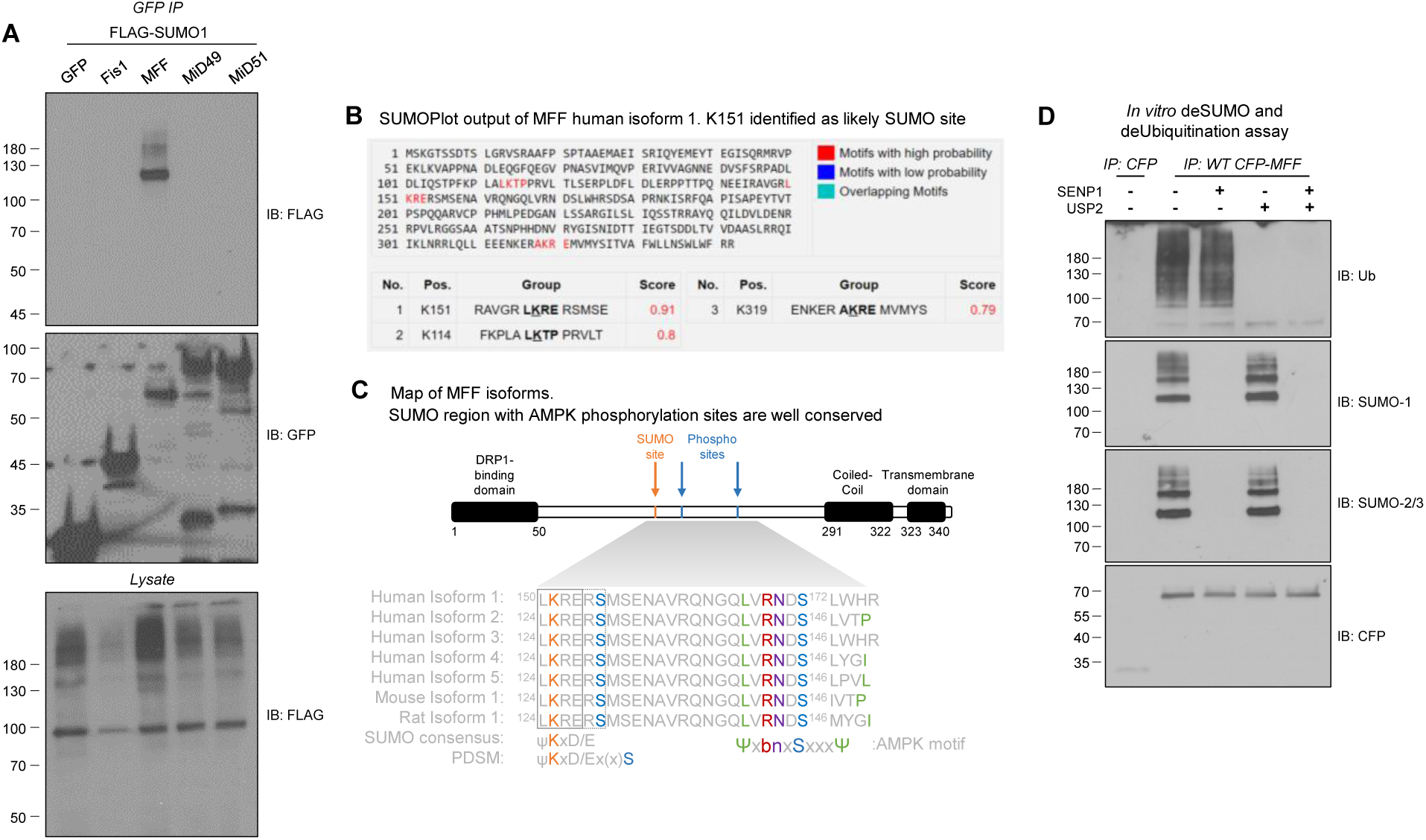
The MFF SUMOylation sequence is well conserved, lies within a PDSM motif, and SUMOylation is independent of MFF ubiquitination. **(A)** Screen of GFP-tagged DRP1 receptors for SUMOylation. HEK293T cells were co-transfected with GFP-Fis1 (rat) GFP-MFF (isoform 1, human), MiD49-GFP or MiD51-GFP (human) and FLAG-SUMO1. GFP-TRAP was used to IP GFP-tagged proteins and immunoprecipitates were blotted for FLAG and GFP. **(B)** SUMOplot^TM^ output for SUMO consensus motifs within the human MFF isoform 1 protein sequence. **(C)** Alignment of human, mouse and rat isoforms of MFF. Schematic representation of MFF, showing the N-terminal DRP1 binding domain, SUMO consensus sequence at ^150^LKRE^153^ (ψKxD/E, where ψ=hydrophobic amino acid, x=any amino acid) and the phosphorylation sites at Ser^155^ and Ser^172^, the coiled-coil domain towards the C-terminus, and the single transmembrane domain at the extreme C-terminus. The LKRE motif, as well as the two phosphorylation sites, are conserved among human isoforms 1-5, and the mouse and rat sequences. Due to alternate splicing of human MFF, human isoform 1 has phosphorylation sites at Ser^155^ and Ser^172^, whereas these correspond to Ser^129^ and Ser^146^ in the other sequences. PDSM=phosphorylation-dependent SUMO consensus motif. The alternative splicing results in a different amino acid sequence C-terminally to the AMPK site at Ser^172^. All isoforms contain many of the elements of the AMPK motif ΨxbnxSxxxΨ (Ψ=hydrophobic amino acid, b=basic amino acid, n=neutral amino acid) (Hardie et al., 2016). Isoforms correspond to Uniprot entries. **(D)** *In vitro* deSUMOylation and deubiquitination assay of MFF. CFP-MFF (WT) from transfected HEK293T cells was isolated on GFP-TRAP beads. The beads were equally separated into different tubes and treated with 100nM SENP1, 500nM USP2 (or both) for 2hrs at 37°C. Samples were resolved by SDS-PAGE and probed for SUMO1, SUMO2/3 and ubiquitin.

**Figure S2.**
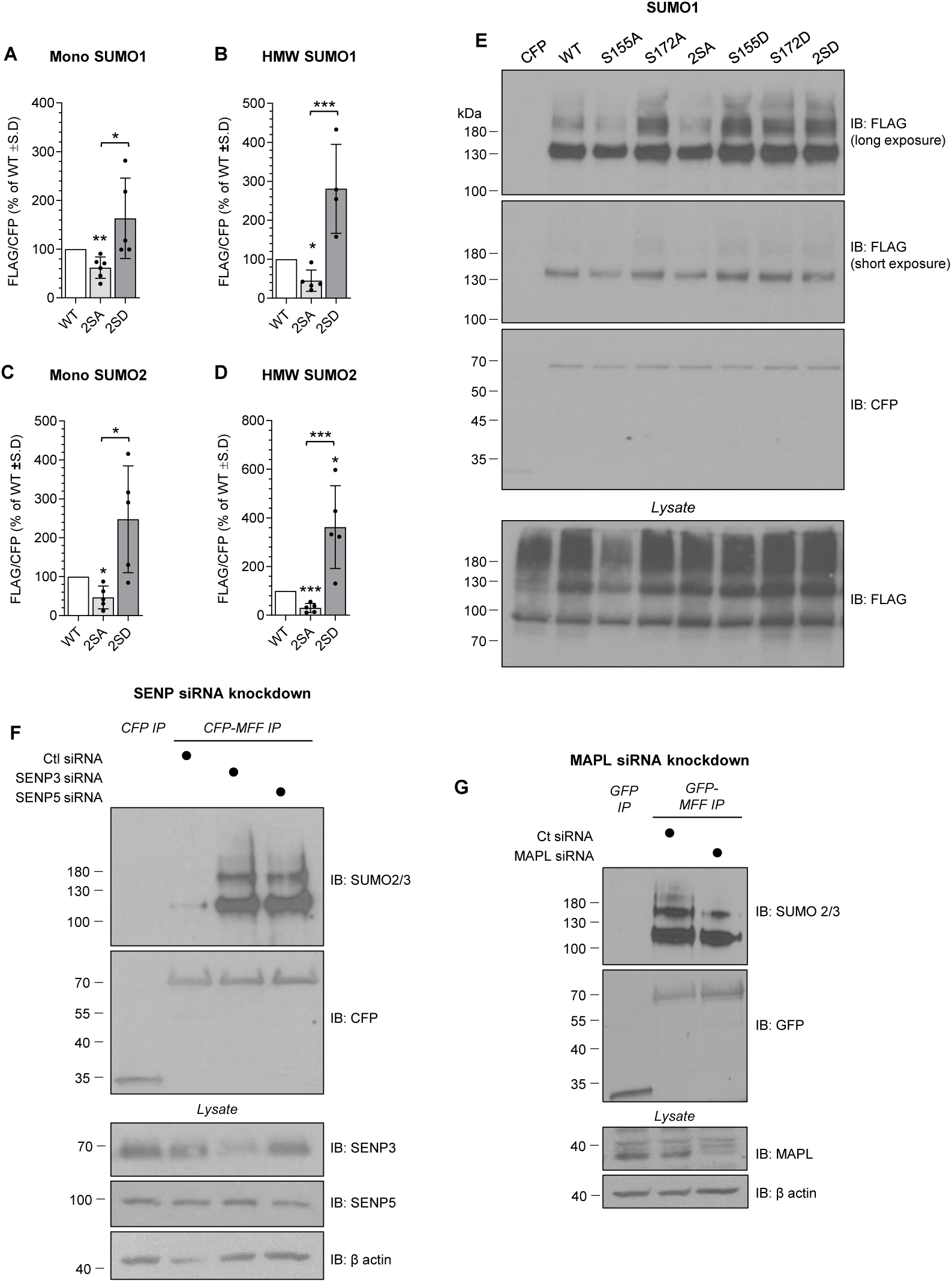
The SUMOylation status of MFF is regulated by AMPK phosphorylation, MAPL, SENP3 and SENP5. (**A-D**) Independent quantification of the mono-SUMO band and the higher molecular weight (HMW) bands of SUMOylated MFF. Corresponds to Fig 3A-D. The mono-SUMO band at 130kDa (**A**) and the higher molecular weight bands (**B**) were quantified, normalised to the CFP blot and expressed as mean percentage of wild-type MFF. (**C-D**) As in **A-B** but corresponds to SUMO2. Data generated from 4 (**B**) or 5 (**A, C-D**) independent experiments. One sample t-test used to determine significance between 2SA/D and WT, two sample t-test used to determine significance between 2SA and 2SD. p*<0.05, p**<0.01, p***<0.005. (**E**) SUMOylation of MFF Ser^155^ and Ser^172^ phospho-mutants (showing full blot from Fig 3A). HEK293T cells were co-transfected with the indicated CFP-MFF mutants and FLAG-SUMO1. GFP-TRAP performed on lysate, resolved by SDS-PAGE and probed for FLAG and CFP. (**F**) SENP3/5 deSUMOylates MFF. HEK293T cells were co-transfected with CFP-MFF (WT) and siRNA targeted against SENP3 or SENP5 (100nM, 48hrs). Immunoprecipitates were probed for SUMO2/3. Lysates probed for SENP3, SENP5 and β-actin. (**G**) MAPL (also called Mul1) is an E3 ligase of MFF. HEK293T were co-transfected with GFP-MFF (WT) and MAPL siRNA at 20nM for 48hrs. Immunoprecipitates were probed for SUMO2/3 and lysate probed for MAPL and β-actin.

**Figure S3.**
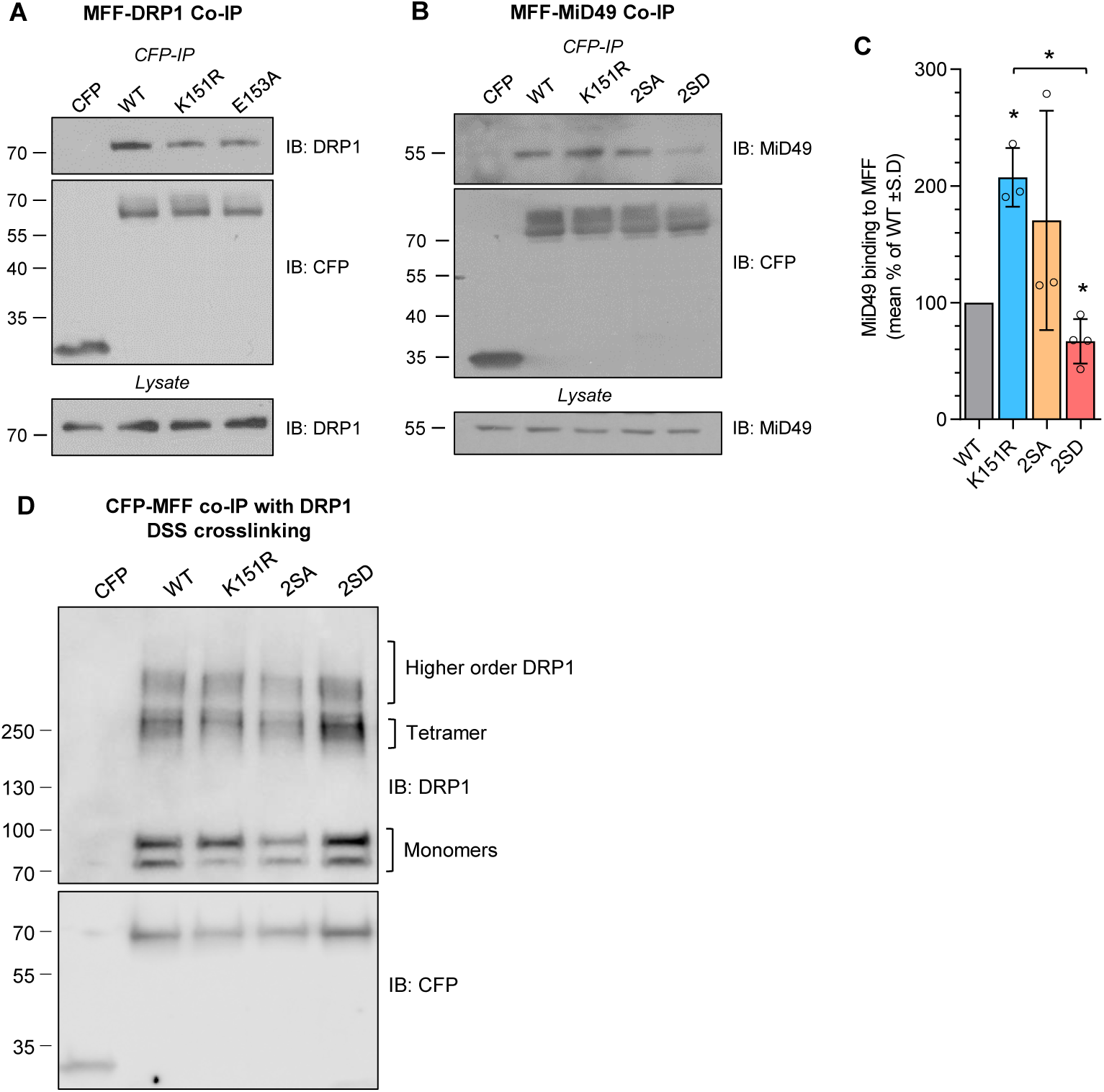
Post-translational modification of MFF alters endogenous DRP1 and MiD49 binding. (**A**) Western blot of endogenous DRP1 binding to MFF SUMO mutants. HEK293T cells were transfected with either CFP-MFF (WT, K151R or E153A), co-IP carried out on lysate using GFP-TRAP and samples immunoblotted for endogenous DRP1. (**B**) Western blot of endogenous MiD49 binding to MFF mutants. HEK293T cells were co-transfected with the indicated CFP-MFF constructs. GFP-TRAP was performed to isolate CFP-MFF, resolved by SDS-PAGE and Western blotted for MiD49. (**C**) Quantification of MiD49 binding to MFF mutants. MiD49 signal was normalised to the CFP reprobe and represented as a percentage of WT±S.D. Data generated from 3/4 independent experiments, one sample t-test used to determine significance from WT, one-way ANOVA used between groups, p*<0.05. (**D**) Proteins were chemically crosslinked with 1mM DSS for 30 minutes prior to lysis. Cells were then lysed on ice and co-IP performed on samples as in figure 4. Samples were resolved on 4-20% gradient gels and probed for DRP1. No difference observed between mutants in the ratio of oligomeric/monomer ratio.

**Figure S4.**
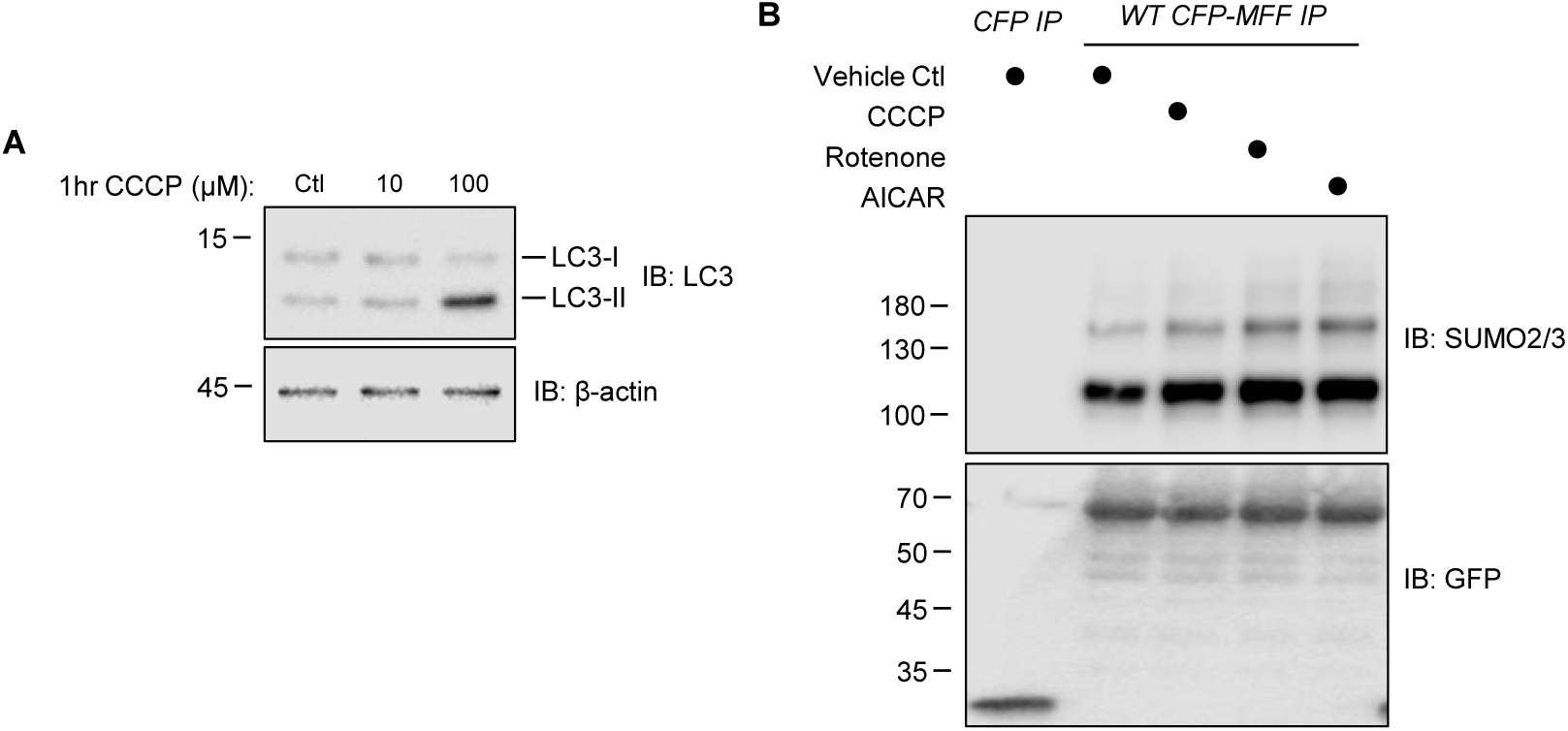
Mitochondrial stressors and AMPK activation promote MFF SUMOylation. CCCP does not activate mitophagy under our conditions. **(A)** Western blot of HEK293T cells treated with either 10 or 100µM CCCP for 1hr (DMSO as vehicle control). Cells were lysed in Laemmli buffer, boiled and resolved by SDS-PAGE. Blots were probed using LC3A/B and β-actin. Appearance of lower LC3-II band indicates formation of autophagosomes. **(B)** Western blot of CFP-MFF WT IP from transfected HEK293T treated with CCCP (10µM, 1hr), Rotenone (250ng/mL, 1hr) or AICAR (1mM, 1hr), and immunoprecipitates probed for SUMO2/3. Corresponds to blot in Fig 5F.

**Figure S5.**
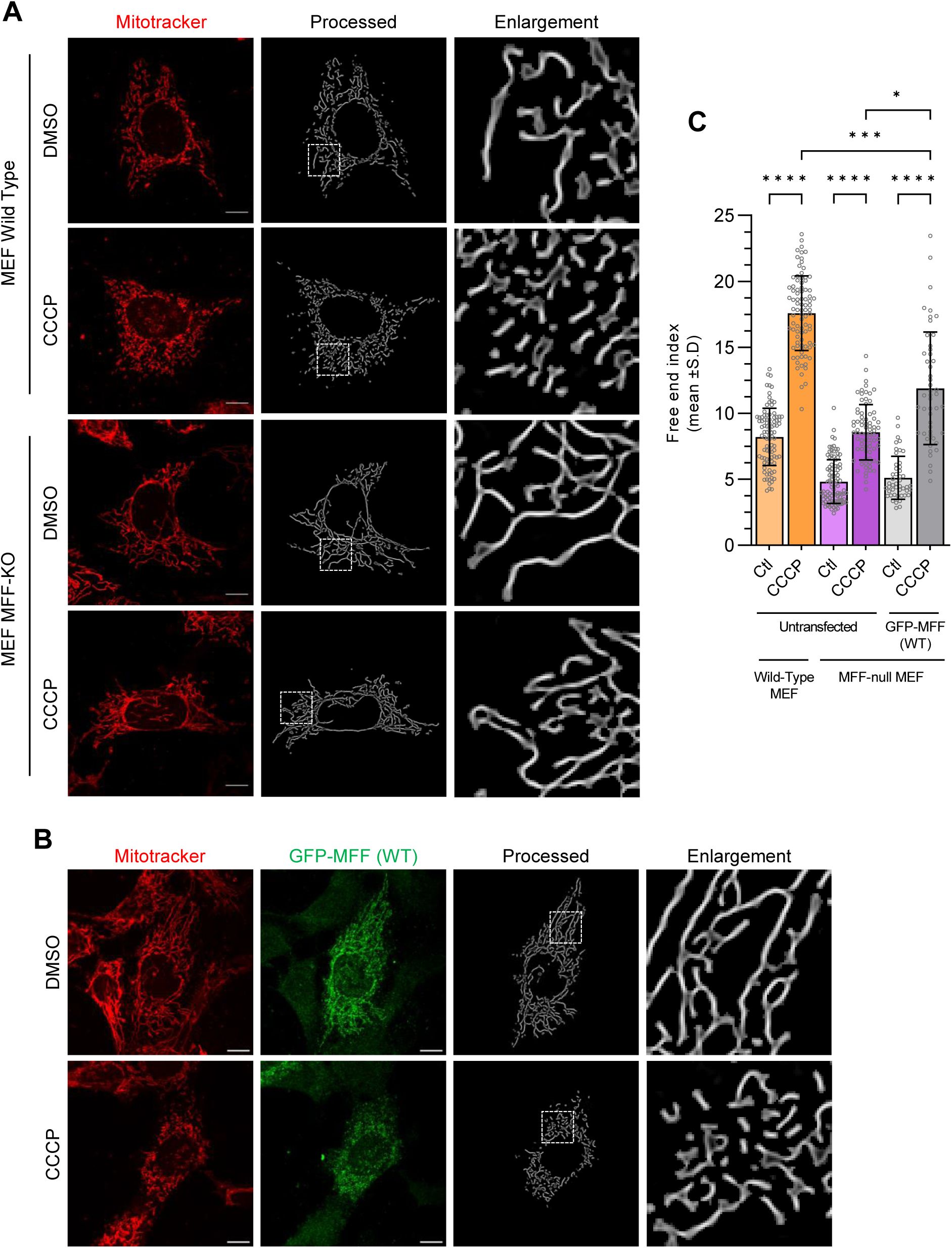
Fragmentation of MEF wild-type and MFF-KO cells with CCCP. **(A)** Confocal imaging of MEF wild-type and MFF-KO cells treated with CCCP to induce fragmentation. Cells were pretreated with mitotracker before application of 10µM CCCP for 1hr. Confocal images were processed (as described in methods) to generate an outline of the mitochondrial network. Scale bar 10µm. Enlargements show zoomed region of highlighted area. **(B)** GFP-MFF (WT) expression in MEF MFF-KO cells promotes fragmentation following CCCP treatment. Using lenti virus to express GFP-MFF in the MFF-KO MEF cells, mitochondria were stained with mitotracker deep red, and treated with CCCP (10µM, 1hr) before fixing. Red channel shows mitochondrial stain, green channel shows GFP stain to confirm GFP-MFF expression. Scale bar 10µm, enlargements show zoomed region of highlighted area. **(C)** Quantification of the extent of fragmentation (mean free end index) following CCCP treatment of MEF wild-type and MFF-KO cells. Data generated form three independent experiments, 82-95 cells imaged (wild-type and MFF-KO MEF cells), data generated from two independent experiments, 47-49 cells imaged for GFP-MFF (WT) expression. Kruskal-Wallis test, p*<0.05, p***<0.001, p****<0.0001.

## Author contributions

Conceptualisation: RS, KAW, JMH

Methodology: RS, KAW, SC

Experimentation: RS, NSR, KAW

Data analysis: RS, NSR

Supervision: KAW, JMH

Writing – original draft: RS

CG first identified MFF as a SUMO substrate.

SC helped with imaging analysis and writing the ImageJ macros.

All authors contributed to editing the manuscript.

### Acknowledgements

The authors thank Professors David Chan (Cal Tech, US) and Michael Schrader (Exeter, UK) for supplying the MFF-KO MEF cells. We thank Dr. Paul Murphy (Professor Ron Hay lab, Dundee, UK) for advice and supplying the USP2 for the *in vitro* deubiquitination assay. The authors gratefully acknowledge the Wolfson Bioimaging Facility (Bristol, UK) for their support and assistance in this work

**Funding:** This work was supported by the BBSRC (BB/R00787X/1), Wellcome Trust (105384/Z/14/A) and Leverhulme Trust (RPG-2019-191). NSR is funded by a Marshall Scholarship.

**Competing interests:** Authors declare that they have no competing interests.

**Data and materials availability:** All data are available in the main text or the supplementary materials.

## References

1. de Brito, O.M., and L. Scorrano. 2008. Mitofusin 2 tethers endoplasmic reticulum to mitochondria. Nature. 456:605–610. doi:10.1038/nature07534.

2. Cagalinec, M., D. Safiulina, M. Liiv, J. Liiv, V. Choubey, P. Wareski, V. Veksler, and A. Kaasik. 2013. Principles of the mitochondrial fusion and fission cycle in neurons. J. Cell Sci. 126:2187–2197. doi:10.1242/jcs.118844.

3. Chan, D.C. 2020. Mitochondrial Dynamics and Its Involvement in Disease. Annu. Rev. Pathol. Mech. Dis. 15:235–259. doi:10.1146/annurev-pathmechdis-012419-032711.

4. Chen, H., A. Chomyn, and D.C. Chan. 2005. Disruption of Fusion Results in Mitochondrial Heterogeneity and Dysfunction. J. Biol. Chem. 280:26185–26192. doi:10.1074/JBC.M503062200.

5. Detmer, S.A., and D.C. Chan. 2007. Functions and dysfunctions of mitochondrial dynamics. Nat. Rev. Mol. Cell Biol. 8:870–879. doi:10.1038/nrm2275.

6. Drake, J.C., R.J. Wilson, R.C. Laker, Y. Guan, H.R. Spaulding, A.S. Nichenko, W. Shen, H. Shang, M. V. Dorn, K. Huang, M. Zhang, A.B. Bandara, M.H. Brisendine, J.A. Kashatus, P.R. Sharma, A. Young, J. Gautam, R. Cao, H. Wallrabe, P.A. Chang, M. Wong, E.M. Desjardins, S.A. Hawley, G.J. Christ, D.F. Kashatus, C.L. Miller, M.J. Wolf, A. Periasamy, G.R. Steinberg, D.G. Hardie, and Z. Yan. 2021. Mitochondria-localized AMPK responds to local energetics and contributes to exercise and energetic stress-induced mitophagy. Proc. Natl. Acad. Sci. U. S. A. 118:e2025932118. doi:10.1073/pnas.2025932118.

7. Ducommun, S., M. Deak, D. Sumpton, R.J. Ford, A. Núñez Galindo, M. Kussmann, B. Viollet, G.R. Steinberg, M. Foretz, L. Dayon, N.A. Morrice, and K. Sakamoto. 2015. Motif affinity and mass spectrometry proteomic approach for the discovery of cellular AMPK targets: Identification of mitochondrial fission factor as a new AMPK substrate. Cell. Signal. 27:978–988. doi:10.1016/J.CELLSIG.2015.02.008.

8. Erker, Y., H. Neyret-Kahn, J.S. Seeler, A. Dejean, A. Atfi, and L. Levy. 2013. Arkadia, a novel SUMO-targeted ubiquitin ligase involved in PML degradation. Mol. Cell. Biol. 33:2163–2177. doi:10.1128/MCB.01019-12.

9. Flotho, A., and F. Melchior. 2013. Sumoylation: a regulatory protein modification in health and disease. Annu. Rev. Biochem. 82:357–385. doi:10.1146/annurev-biochem-061909-093311.

10. Frank, S., B. Gaume, E.S. Bergmann-Leitner, W.W. Leitner, E.G. Robert, F. Catez, C.L. Smith, and R.J. Youle. 2001. The Role of Dynamin-Related Protein 1, a Mediator of Mitochondrial Fission, in Apoptosis. Dev. Cell. 1:515–525. doi:10.1016/S1534-5807(01)00055-7.

11. Fu, C., K. Ahmed, H. Ding, X. Ding, J. Lan, Z. Yang, Y. Miao, Y. Zhu, Y. Shi, J. Zhu, H. Huang, and X. Yao. 2005. Stabilization of PML nuclear localization by conjugation and oligomerization of SUMO-3. Oncogene. 24:5401–5413. doi:10.1038/sj.onc.1208714.

12. Gandre-Babbe, S., and A.M. van der Bliek. 2008. The Novel Tail-anchored Membrane Protein Mff Controls Mitochondrial and Peroxisomal Fission in Mammalian Cells. Mol. Biol. Cell. 19:2402–2412. doi:10.1091/mbc.e07-12-1287.

13. Gomes, L.C., G. Di Benedetto, and L. Scorrano. 2011. During autophagy mitochondria elongate, are spared from degradation and sustain cell viability. Nat. Cell Biol. 13:589–598. doi:10.1038/ncb2220.

14. Guo, C., K.L. Hildick, J. Luo, L. Dearden, K.A. Wilkinson, and J.M. Henley. 2013. SENP3-mediated deSUMOylation of dynamin-related protein 1 promotes cell death following ischaemia. EMBO J. 32:1514–1528. doi:10.1038/emboj.2013.65.

15. Guo, C., K.A. Wilkinson, A.J. Evans, P.P. Rubin, and J.M. Henley. 2017. SENP3-mediated deSUMOylation of Drp1 facilitates interaction with Mff to promote cell death. Sci. Rep. 7:43811. doi:10.1038/srep43811.

16. Harder, Z., R. Zunino, and H. McBride. 2004. Sumo1 Conjugates Mitochondrial Substrates and Participates in Mitochondrial Fission. Curr. Biol. 14:340–345. doi:10.1016/J.CUB.2004.02.004.

17. Hardie, D.G., B.E. Schaffer, and A. Brunet. 2016. AMPK: An Energy-Sensing Pathway with Multiple Inputs and Outputs. Trends Cell Biol. 26:190–201. doi:10.1016/j.tcb.2015.10.013.

18. Herzig, S., and R.J. Shaw. 2018. AMPK: guardian of metabolism and mitochondrial homeostasis. Nat. Rev. Mol. Cell Biol. 19:121–135. doi:10.1038/nrm.2017.95.

19. Hietakangas, V., J. Anckar, H.A. Blomster, M. Fujimoto, J.J. Palvimo, A. Nakai, and L. Sistonen. 2006. PDSM, a motif for phosphorylation-dependent SUMO modification. Proc. Natl. Acad. Sci. U. S. A. 103:45–50. doi:10.1073/pnas.0503698102.

20. Iwasawa, R., A.-L. Mahul-Mellier, C. Datler, E. Pazarentzos, and S. Grimm. 2011. Fis1 and Bap31 bridge the mitochondria–ER interface to establish a platform for apoptosis induction. EMBO J. 30:556–558. doi:10.1038/EMBOJ.2010.346.

21. Kageyama, Y., M. Hoshijima, K. Seo, D. Bedja, P. Sysa-Shah, S.A. Andrabi, W. Chen, A. Höke, V.L. Dawson, T.M. Dawson, K. Gabrielson, D.A. Kass, M. Iijima, and H. Sesaki. 2014. Parkin-independent mitophagy requires Drp1 and maintains the integrity of mammalian heart and brain. EMBO J. 33:2798–2813. doi:10.15252/embj.201488658.

22. Kageyama, Y., Z. Zhang, R. Roda, M. Fukaya, J. Wakabayashi, N. Wakabayashi, T.W. Kensler, P.H. Reddy, M. Iijima, and H. Sesaki. 2012. Mitochondrial division ensures the survival of postmitotic neurons by suppressing oxidative damage. J. Cell Biol. 197:535–551. doi:10.1083/jcb.201110034.

23. Kleele, T., T. Rey, J. Winter, S. Zaganelli, D. Mahecic, H.P. Lambert, F.P. Ruberto, M. Nemir, T. Wai, T. Pedrazzini, and S. Manley. 2021. Distinct fission signatures predict mitochondrial degradation or biogenesis. Nat. 2021 5937859. 593:435–439. doi:10.1038/s41586-021-03510-6.

24. Lallemand-Breitenbach, V., M. Jeanne, S. Benhenda, R. Nasr, M. Lei, L. Peres, J. Zhou, J. Zhu, B. Raught, and H. de Thé. 2008. Arsenic degrades PML or PML– RARα through a SUMO-triggered RNF4/ubiquitin-mediated pathway. Nat. Cell Biol. 10:547–555. doi:10.1038/ncb1717.

25. Lallemand-Breitenbach, V., and H. de Thé. 2010. PML nuclear bodies. Cold Spring Harb. Perspect. Biol. 2:a000661. doi:10.1101/cshperspect.a000661.

26. Leboucher, G.P., Y.C. Tsai, M. Yang, K.C. Shaw, M. Zhou, T.D. Veenstra, M.H. Glickman, and A.M. Weissman. 2012. Stress-Induced Phosphorylation and Proteasomal Degradation of Mitofusin 2 Facilitates Mitochondrial Fragmentation and Apoptosis. Mol. Cell. 47:547–557. doi:10.1016/j.molcel.2012.05.041.

27. Lee, L., R. Seager, Y. Nakamura, K.A. Wilkinson, and J.M. Henley. 2019. Parkin-mediated ubiquitination contributes to the constitutive turnover of mitochondrial fission factor (Mff). PLoS One. 14:e0213116. doi:10.1371/journal.pone.0213116.

28. Lewis, T.L., S.K. Kwon, A. Lee, R. Shaw, and F. Polleux. 2018. MFF-dependent mitochondrial fission regulates presynaptic release and axon branching by limiting axonal mitochondria size. Nat. Commun. 9:5008. doi:10.1038/s41467-018-07416-2.

29. Liu, R., and D.C. Chan. 2015. The mitochondrial fission receptor Mff selectively recruits oligomerized Drp1. Mol. Biol. Cell. 26:4466–4477. doi:10.1091/mbc.E15-08-0591.

30. Losón, O.C., Z. Song, H. Chen, and D.C. Chan. 2013. Fis1, Mff, MiD49, and MiD51 mediate Drp1 recruitment in mitochondrial fission. Mol. Biol. Cell. 24:659–667. doi:10.1091/mbc.e12-10-0721.

31. Matic, I., M. van Hagen, J. Schimmel, B. Macek, S.C. Ogg, M.H. Tatham, R.T. Hay, A.I. Lamond, M. Mann, and A.C.O. Vertegaal. 2008. In vivo identification of human small ubiquitin-like modifier polymerization sites by high accuracy mass spectrometry and an in vitro to in vivo strategy. Mol. Cell. Proteomics. 7:132–144. doi:10.1074/mcp.M700173-MCP200.

32. Mears, J.A., L.L. Lackner, S. Fang, E. Ingerman, J. Nunnari, and J.E. Hinshaw. 2011. Conformational changes in Dnm1 support a contractile mechanism for mitochondrial fission. Nat. Struct. Mol. Biol. 18:20–27. doi:10.1038/nsmb.1949.

33. Nakada, K., K. Inoue, T. Ono, K. Isobe, A. Ogura, Y.-I. Goto, I. Nonaka, and J.-I. Hayashi. 2001. Inter-mitochondrial complementation: Mitochondria-specific system preventing mice from expression of disease phenotypes by mutant mtDNA. Nat. Med. 7:934–940. doi:10.1038/90976.

34. Osellame, L.D., A.P. Singh, D.A. Stroud, C.S. Palmer, D. Stojanovski, R. Ramachandran, and M.T. Ryan. 2016. Cooperative and independent roles of the Drp1 adaptors Mff, MiD49 and MiD51 in mitochondrial fission. J. Cell Sci. 129:2170–2181. doi:10.1242/jcs.185165.

35. Otera, H., N. Miyata, O. Kuge, and K. Mihara. 2016. Drp1-dependent mitochondrial fission via MiD49/51 is essential for apoptotic cristae remodeling. J. Cell Biol. 212:531–544. doi:10.1083/jcb.201508099.

36. Otera, H., C. Wang, M.M. Cleland, K. Setoguchi, S. Yokota, R.J. Youle, and K. Mihara. 2010. Mff is an essential factor for mitochondrial recruitment of Drp1 during mitochondrial fission in mammalian cells. J. Cell Biol. 191:1141–1158. doi:10.1083/jcb.201007152.

37. Palmer, C.S., K.D. Elgass, R.G. Parton, L.D. Osellame, D. Stojanovski, and M.T. Ryan. 2013. Adaptor proteins MiD49 and MiD51 can act independently of Mff and Fis1 in Drp1 recruitment and are specific for mitochondrial fission. J. Biol. Chem. 288:27584–27593. doi:10.1074/jbc.M113.479873.

38. Palmer, C.S., L.D. Osellame, D. Laine, O.S. Koutsopoulos, A.E. Frazier, and M.T. Ryan. 2011. MiD49 and MiD51, new components of the mitochondrial fission machinery. EMBO Rep. 12:565–573. doi:10.1038/embor.2011.54.

39. Pangou, E., O. Bielska, L. Guerber, S. Schmucker, A. Agote-Arán, T. Ye, Y. Liao, M. Puig-Gamez, E. Grandgirard, C. Kleiss, Y. Liu, E. Compe, Z. Zhang, R. Aebersold, R. Ricci, and I. Sumara. 2021. A PKD-MFF signaling axis couples mitochondrial fission to mitotic progression. Cell Rep. 35:109129. doi:10.1016/j.celrep.2021.109129.

40. Park, Y.Y., O.T.K. Nguyen, H. Kang, and H. Cho. 2014. MARCH5-mediated quality control on acetylated Mfn1 facilitates mitochondrial homeostasis and cell survival. Cell Death Dis. 5:1172–1184. doi:10.1038/cddis.2014.142.

41. Pernas, L., and L. Scorrano. 2016. Mito-Morphosis: Mitochondrial Fusion, Fission, and Cristae Remodeling as Key Mediators of Cellular Function. Annu. Rev. Physiol. 78:505–531. doi:10.1146/annurev-physiol-021115-105011.

42. Prudent, J., R. Zunino, A. Sugiura, S. Mattie, G.C. Shore, and H.M. McBride. 2015. MAPL SUMOylation of Drp1 Stabilizes an ER/Mitochondrial Platform Required for Cell Death. Mol. Cell. 59:941–955. doi:10.1016/J.MOLCEL.2015.08.001.

43. Qi, X., M.H. Disatnik, N. Shen, R.A. Sobel, and D. Mochly-Rosen. 2011. Aberrant mitochondrial fission in neurons induced by protein kinase Cδ under oxidative stress conditions in vivo. Mol. Biol. Cell. 22:256–265. doi:10.1091/mbc.E10-06-0551.

44. Rambold, A.S., B. Kostelecky, N. Elia, and J. Lippincott-Schwartz. 2011. Tubular network formation protects mitochondria from autophagosomal degradation during nutrient starvation. Proc. Natl. Acad. Sci. 108:10190–10195. doi:10.1073/pnas.1107402108.

45. Rocca, D.L., K.A. Wilkinson, and J.M. Henley. 2017. SUMOylation of FOXP1 regulates transcriptional repression via CtBP1 to drive dendritic morphogenesis. Sci. Rep. 7:877. doi:10.1038/s41598-017-00707-6.

46. Seabright, A.P., N.H.F. Fine, J.P. Barlow, S.O. Lord, I. Musa, A. Gray, J.A. Bryant, M. Banzhaf, G.G. Lavery, D.G. Hardie, D.J. Hodson, A. Philp, and Y.C. Lai. 2020. AMPK activation induces mitophagy and promotes mitochondrial fission while activating TBK1 in a PINK1-Parkin independent manner. FASEB J. 34:6284–6301. doi:10.1096/fj.201903051R.

47. Shen, Q., K. Yamano, B.P. Head, S. Kawajiri, J.T.M. Cheung, C. Wang, J.H. Cho, N. Hattori, R.J. Youle, and A.M. Van Der Bliek. 2014. Mutations in Fis1 disrupt orderly disposal of defective mitochondria. Mol. Biol. Cell. 25:145–159. doi:10.1091/mbc.E13-09-0525.

48. Smirnova, E., L. Griparic, D.L. Shurland, and A.M. Van der Bliek. 2001. Dynamin-related protein Drp1 is required for mitochondrial division in mammalian cells. Mol. Biol. Cell. 12:2245–2256. doi:10.1091/mbc.12.8.2245.

49. Sriramachandran, A.M., K. Meyer-Teschendorf, S. Pabst, H.D. Ulrich, N.H. Gehring, K. Hofmann, G.J.K. Praefcke, and R.J. Dohmen. 2019. Arkadia/RNF111 is a SUMO-targeted ubiquitin ligase with preference for substrates marked with SUMO1-capped SUMO2/3 chain. Nat. Commun. 2019 101. 10:3678. doi:10.1038/s41467-019-11549-3.

50. Taguchi, N., N. Ishihara, A. Jofuku, T. Oka, and K. Mihara. 2007. Mitotic phosphorylation of dynamin-related GTPase Drp1 participates in mitochondrial fission. J. Biol. Chem. 282:11521–11529. doi:10.1074/jbc.M607279200.

51. Tatham, M.H., E. Jaffray, O.A. Vaughan, J.M.P. Desterro, C.H. Botting, J.H. Naismith, and R.T. Hay. 2001. Polymeric Chains of SUMO-2 and SUMO-3 are Conjugated to Protein Substrates by SAE1/SAE2 and Ubc9. J. Biol. Chem. 276:35368–35374. doi:10.1074/jbc.M104214200.

52. Tilokani, L., S. Nagashima, V. Paupe, and J. Prudent. 2018. Mitochondrial dynamics: overview of molecular mechanisms. Essays Biochem. 62:341–360. doi:10.1042/EBC20170104.

53. Tondera, D., S. Grandemange, A. Jourdain, M. Karbowski, Y. Mattenberger, S. Herzig, S. Da Cruz, P. Clerc, I. Raschke, C. Merkwirth, S. Ehses, F. Krause, D.C. Chan, C. Alexander, C. Bauer, R. Youle, T. Langer, and J.-C. Martinou. 2009. SLP-2 is required for stress-induced mitochondrial hyperfusion. EMBO J. 28:1589–1600. doi:10.1038/EMBOJ.2009.89.

54. Toyama, E.Q., S. Herzig, J. Courchet, T.L. Lewis, O.C. Loson, K. Hellberg, N.P. Young, H. Chen, F. Polleux, D.C. Chan, and R.J. Shaw. 2016. AMP-activated protein kinase mediates mitochondrial fission in response to energy stress. Science. 351:275–281. doi:10.1126/science.aab4138.

55. Twig, G., A. Elorza, A.J.A. Molina, H. Mohamed, J.D. Wikstrom, G. Walzer, L. Stiles, S.E. Haigh, S. Katz, G. Las, J. Alroy, M. Wu, B.F. Py, J. Yuan, J.T. Deeney, B.E. Corkey, and O.S. Shirihai. 2008. Fission and selective fusion govern mitochondrial segregation and elimination by autophagy. EMBO J. 27:433–446. doi:10.1038/sj.emboj.7601963.

56. Valente, A.J., L.A. Maddalena, E.L. Robb, F. Moradi, and J.A. Stuart. 2017. A simple ImageJ macro tool for analyzing mitochondrial network morphology in mammalian cell culture. Acta Histochem. 3:315–326. doi:10.1016/j.acthis.2017.03.001.

57. Wai, T., and T. Langer. 2016. Mitochondrial Dynamics and Metabolic Regulation. Trends Endocrinol. Metab. 27:105–117. doi:10.1016/j.tem.2015.12.001.

58. Waters, E., K.A. Wilkinson, A.L. Harding, R.E. Carmichael, D. Robinson, H.E. Colley, and C. Guo. 2022. The SUMO protease SENP3 regulates mitochondrial autophagy mediated by Fis1. EMBO Rep. 23:e48754. doi:10.15252/embr.201948754.

59. Wilkinson, K.A., and J.M. Henley. 2010. Mechanisms, regulation and consequences of protein SUMOylation. Biochem. J. 428:133–145. doi:10.1042/BJ20100158.

60. Yu, R., S.-B. Jin, M. Ankarcrona, U. Lendahl, M. Nistér, and J. Zhao. 2021. The Molecular Assembly State of Drp1 Controls its Association With the Mitochondrial Recruitment Receptors Mff and MIEF1/2. Front. Cell Dev. Biol. 9:706687. doi:10.3389/fcell.2021.706687.

61. Yu, R., T. Liu, S.-B. Jin, C. Ning, U. Lendahl, M. Nistér, and J. Zhao. 2017. MIEF1/2 function as adaptors to recruit Drp1 to mitochondria and regulate the association of Drp1 with Mff. Sci. Rep. 7:880. doi:10.1038/s41598-017-00853-x.

62. Zhao, J., T. Liu, S. Jin, X. Wang, M. Qu, P. Uhlén, N. Tomilin, O. Shupliakov, U. Lendahl, and M. Nistér. 2011. Human MIEF1 recruits Drp1 to mitochondrial outer membranes and promotes mitochondrial fusion rather than fission. EMBO J. 30:2762–2778. doi:10.1038/emboj.2011.198.

63. Zhong, S., S. Müller, S. Ronchetti, P.S. Freemont, A. Dejean, and P.P. Pandolfi. 2000. Role of SUMO-1-modified PML in nuclear body formation. Blood. 95:2748–2752.

64. Zunino, R., E. Braschi, L. Xu, and H.M. McBride. 2009. Translocation of SenP5 from the nucleoli to the mitochondria modulates DRP1-dependent fission during mitosis. J. Biol. Chem. 284:17783–17795. doi:10.1074/jbc.M901902200.

65. Zunino, R., A. Schauss, P. Rippstein, M. Andrade-Navarro, and H.M. McBride. 2007. The SUMO protease SENP5 is required to maintain mitochondrial morphology and function. J. Cell Sci. 120:1178–1188. doi:10.1242/jcs.03418.

